# Meta-analysis of Gene Expression Microarray Datasets in Chronic Obstructive Pulmonary Disease

**DOI:** 10.1101/671206

**Authors:** Lavida R. K. Rogers, Madison Verlinde, George I. Mias

## Abstract

Chronic obstructive pulmonary disease (COPD) was classified by the Centers for Disease Control and Prevention in 2014 as the 3^rd^ leading cause of death in the United States (US). The main cause of COPD is exposure to tobacco smoke and air pollutants. Problems associated with COPD include under-diagnosis of the disease and an increase in the number of smokers worldwide. The goal of our study is to identify disease variability in the gene expression profiles of COPD subjects compared to controls. We used pre-existing, publicly available microarray expression datasets to conduct a meta-analysis. Our inclusion criteria for microarray datasets selected for smoking status, age and sex of blood donors reported. Our datasets used Affymetrix, Agilent microarray platforms (7 datasets, 1,262 samples). We re-analyzed the curated raw microarray expression data using R packages, and used Box-Cox power transformations to normalize datasets. To identify significant differentially expressed genes we ran an analysis of variance with a linear model with disease state, age, sex, smoking status and study as effects that also included binary interactions. We found 1,513 statistically significant (Benjamini-Hochberg-adjusted p-value <0.05) differentially expressed genes with respect to disease state (COPD or control). We further filtered these genes for biological effect using results from a Tukey test post-hoc analysis (Benjamini-Hochberg-adjusted p-value <0.05 and 10% two-tailed quantiles of mean differences between COPD and control), to identify 304 genes. Through analysis of disease, sex, age, and also smoking status and disease interactions we identified differentially expressed genes involved in a variety of immune responses and cell processes in COPD. We also trained a logistic regression model using the 304 genes as features, which enabled prediction of disease status with 84% accuracy. Our results give potential for improving the diagnosis of COPD through blood and highlight novel gene expression disease signatures.

## Introduction

Chronic obstructive pulmonary disease (COPD) impairs lung function and reduces lung capacity. In COPD there is inflammation of the bronchial tubes (chronic bronchitis) [1] and destruction of the air sacs (emphysema) [2] within the lungs [3–6]. Chronic bronchitis and emphysema often occur together and are grouped under COPD [1, 2]. Furthermore, the Global Initiative for Chronic Obstructive Lung Disease (GOLD) describes COPD as a common and preventable disease that is caused by exposure to harmful particles and gases that affect the airways and alveolar of the lungs [7, 8]. Individuals with COPD experience shortness of breath due to lowered concentrations of oxygen in the blood and a chronic cough accompanied by mucus production [1–4, 6]. COPD progresses with time and the damage caused to the lungs is irreversible [8, 9]. However, there are treatments available to control disease progression [8, 9].

COPD, the 3^rd^ leading cause of death in the United States (US), is expected to rise in 15 years to the leading cause of death [8–10]. Globally, there were over 250 million cases of COPD reported in 2016 and in 2015 3.17 million individuals died from the disease [5]. COPD is prevalent in low- and middle-income countries with over 90% of COPD cases occurring in these areas [5, 10]. The disease is mainly caused by tobacco exposure through smoking cigarettes or second-hand exposure to smoke [8, 9]. In addition to this, continuous exposure to other irritants such as burning fuels, chemicals, polluted air and dust can lead to COPD [5]. Cigarette smoke exposes the lungs to large amounts of oxidants that induce inflammation of the airways. Previous research on bronchial biopsies highlighted the presence of increased concentrations of inflammatory cells throughout the lungs [11, 12]. Studies have also suggested that COPD acts like an autoimmune disease due to persistent inflammation even after smoking has ceased [12–14]. In addition to environmental pollutants, there is also also a genetic deficiency, alpha-1 antitrypsin deficiency (AATD), that is associated with COPD [8]. AATD protects the lungs, and without it the lungs become vulnerable to COPD. The prevalence of COPD is expected to rise due to increasing smoking rates and larger populations of elderly individuals in many countries [5].

COPD is often underdiagnosed and despite tobacco exposure being the highest risk factor, not all smokers get COPD, and non-smokers can also develop COPD. Previous work has been done to identify biomarkers for earlier diagnosis of COPD in blood, a non-invasive approach. Bahr et al., compared expression profiles of smokers with COPD and smokers without COPD [15]. They used multiple linear regression to identify candidate genes and pathways. Their results highlighted pathways involved in the immune system and inflammatory response [15]. Another study of blood gene expression in COPD explored using pre-existing gene interaction networks to perform unsupervised clustering to identify COPD disease sub-types [16]. More recently, Reinhold et al., took a different approach by conducting a meta-analysis that identified groups of genes associated with COPD by using consensus modules of gene co-expression. They built networks of genes that were co-expressed and associated with COPD phenotypes [17].

In our meta-analysis, the objective was to identify the effects of age, sex, and smoking status on gene expression in COPD. We investigated gene expression changes in blood for 1,262 samples (574 healthy samples and 688 COPD samples) to identify genes and their associated pathways in COPD. Our study is the largest meta-analysis on blood expression for COPD to date, to the best of our knowledge, and our results offer prospective gene and pathway associations that may be targeted for improving COPD diagnosis and treatment. Our meta-analysis also highlighted disease genes that interact with smoking status, and these genes can be used to further characterize the effects of smoking on COPD development.

## Materials and Methods

We used seven publicly available COPD microarray gene expression datasets in our meta-analysis to evaluate variation in gene expression across samples due to disease status, sex, age and smoking status (Table 1). The 7 expression datasets were from 3 different microarray platforms: Affymetrix GeneChip Human Genome U133 Plus 2.0, Affymetrix Human Gene 1.1 ST Array and Agilent Whole Human Genome Microarray 4×44K. Our current meta-analysis pipeline (similar to Brooks et al. [18]), included 5 main steps (Fig 1): (1) data curation; (2) pre-processing of raw expression data; (3) analysis of variance (ANOVA) on our linear model which compared gene expression changes due to disease state, smoking status, sex and age group; (4) post-hoc analysis using Tukey Honest Significance Difference test (TukeyHSD) for biological significance; and (5) Gene ontology (GO) and pathway enrichment analysis of the differentially expressed and biologically significant genes.

**Table 1.** Description of datasets used in the meta-analysis.

| Database Repository | Dataset Accession | Control | COPD | Platform |
| --- | --- | --- | --- | --- |
| Array Express | E-MTAB-5278 | 181 | 53 | Affymetrix Human Genome Plus 2.0 |
| Array Express | E-MTAB-5279 | 89 | 0 | Affymetrix Human Genome Plus 2.0 |
| GEO | GSE42057 | 42 | 94 | Affymetrix Human Genome Plus 2.0 |
| GEO | GSE47415 | 48 | 0 | Agilent-014850 Whole Human Genome Microarray 4x44K |
| GEO | GSE54837 | 90 | 136 | Affymetrix Human Genome Plus 2.0 |
| GEO | GSE71220 | 44 | 405 | Affymetrix Human Gene 1.1 ST Array |
| GEO | GSE87072 | 80 | 0 | Affymetrix Human Genome Plus 2.0 |

**Fig 1.**
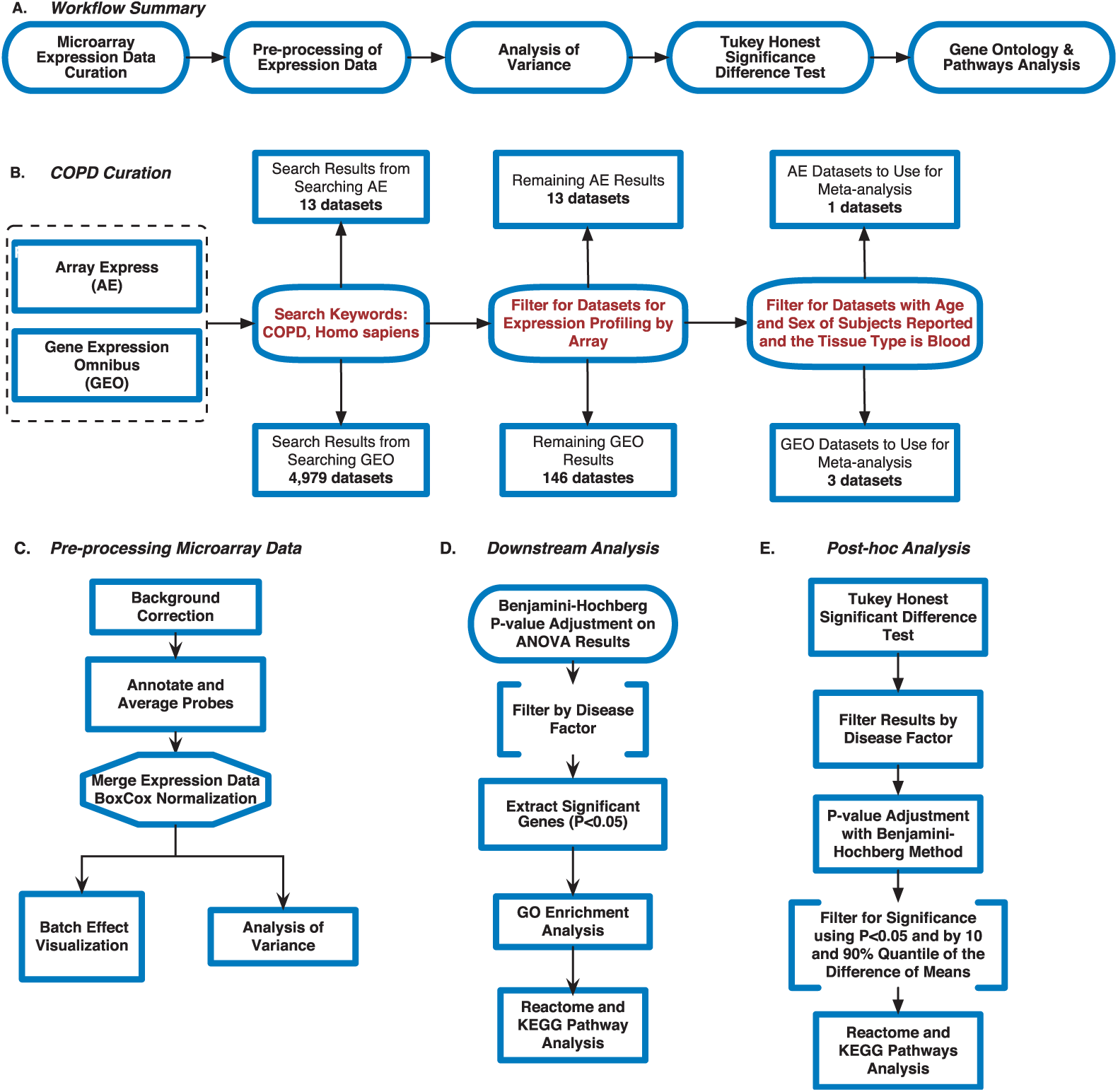
Meta-analysis pipeline for Chronic Obstructive Pulmonary Disease. (A)Summary of workflow used for the meta-analysis,(B) Microarray curation and filtering steps, (C) Pre-processing steps used on the microarray data,(D) Data analysis post ANOVA, (E) post-hoc analysis steps using ANOVA results.

### Microarray Data Curation from Gene Expression Omnibus and Array Express

To gather the datasets for our meta-analysis, we searched the National Center for Biotechnology Information (NCBI)’s data repository, Gene Expression Omnibus (GEO) [19], and the European Bioinformatics Institute (EMBL-EBI)’s data repository, Array Express (AE) [20] for microarray expression data. We used the following keywords to search the repositories: COPD, *Homo sapiens*, blood (whole blood and peripheral blood mononuclear cells) and expression profiling by array (Fig 1B). The search results were further filtered to include datasets where the age, sex and smoking status of the samples were reported (Fig 1B). We found 3 datasets from GEO (GSE42057 [21], GSE71220 [22], GSE54837 [23]) and 1 from AE (E-MTAB-5278 [24]) that met our search criteria (Table 1 and Fig 1B). We conducted an additional search on GEO and AE to find healthy subjects with their smoking history reported to balance our control subjects with our COPD subjects. The search keywords included: *Homo sapiens*, blood, smoking and expression profiling by array. We also filtered these search results for datasets that reported the age, sex and smoking status of subjects. With this additional search, we added 3 more datasets: GSE87072 [25], GSE47415 [26], and E-MTAB-5279 [24] which helped improve the balance between COPD and control subjects (Table 1 and DF1 of online data files).

After selecting the datasets for our meta-analysis, we retrieved the raw microarray expression data for each dataset, and created a demographics file per study, which included sample characteristics using e-utils in Mathematica [27] (Table 2). The demographics files were further filtered to eliminate samples that did not fit our inclusion criteria. For example, GSE71220 included subjects that were using statin drugs [22], and hence we excluded all samples that were receiving treatment from our analysis. For GSE87072, we removed the samples that were moist snuff consumers [25] and only used smokers and non-smokers in our analysis. In our additional search for controls with smoking status reported, we filtered the selected datasets (GSE87072, GSE47415 and E-MTAB-5279) and only used the healthy samples for our analysis. In addition to this, we excluded the subjects in GSE23515 [28] from our analysis because 22 of the 24 samples are duplicates from GSE47415 [26]. Our demographics files were created to include variables that were reported across all samples (see merged Demographics file DF1 of online supplementary data files) because study annotations had not been uniformly reported in the databases (S1 File).

**Table 2.** Sample Characteristics By Dataset.

| Dataset Accession | Sex(M/F) | Smoking Status (S/NS/FS)* | Age Range |
| --- | --- | --- | --- |
| E-MTAB-5279 | 46/43 | 30/29/30 | 24 - 65 |
| EMTAB5278 | 136/98 | 114/60/60 | 41 - 70 |
| GSE42057 | 74/62 | 35/2/99 | 45 - 80 |
| GSE47415 | 24/24 | 24/24/0 | 20 - 64 |
| GSE54837 | 148/78 | 84/6/136 | 40 - 75 |
| GSE71220 | 285/165 | 91/22/336 | 49 - 75 |
| GSE87072 | 80/0 | 40/40/0 | 35 - 60 |

\*S=smoker, NS=non-smoker, FS= former smoker

### Microarray Pre-processing and BoxCox Normalization

To download the raw microarray expression for each dataset we used Mathematica [29]. All raw expression data files were pre-processed in R [30] using R packages specific to each microarray platform (Fig 1C). For the datasets from the Affymetrix Human Genome Plus 2.0 platform, we used the `affy` package [31] for pre-processing all of the .CEL files. The `oligo` [32] and `affycoretools` [33] packages were used to pre-process the data files from the Affymetrix Human Gene 1.1 ST microarry platform, while the `limma` package [34] was used for the data files from the Agilent Whole Human Genome microarray platform. We performed background correction, normalization, and all probes were annotated and summarized (Fig 1C). For the Affymetrix Human Genome Plus 2.0 expression data files, the `expresso` function was used to pre-process the files with the following parameters: background correction with robust multi-array analysis (RMA), correcting the perfect-match (PM) probes, and ‘avdiff’ to calculate expression values [31]. Subsequently, the `avereps` function from `limma` was used to summarize the probes and remove replicates [34]. The Affymetrix Human Gene 1.1 ST data files were also background corrected using RMA, and the probes were summarized and replicates removed using the `avereps` function. As for the Agilent data files, background correction was performed using the `backgroundCorrect` function with NormExp Background Correction as the method from the `limma` package [35]. The probes for both Affymetrix Human Gene 1.1 ST and Agilent were also summarized and replicates were removed using the `avereps` function from `limma`. Once pre-processing was completed, the 8 datasets (Table 1) were merged by common gene symbols into a single matrix file. Using the `ApplyBoxCoxTransform` function and the `StandardizeExtended` function from the MathIOmica (version 1.1.3) package [27, 36] in Mathematica, we performed a Box-Cox power transformation and data standardization on the merged expression file [37] (Fig 1C and DF2 of online supplementary data files).

### Identifying and Visualizing Batch Effects

Conducting meta-analyses by combining expression datasets across different microarray platforms and research labs/studies introduces batch effects/confounding factors to the data. The batch effects can introduce non-biological variation in the data, which affects the interpretation of the results. In order to visualize variation in the expression data across factors, we conducted principal component analysis (PCA) on the expression data and generated PCA plots (Fig 2 and S1 Fig). As we also previously described [18], the study factor is directly related to the microarray platform type. To address this, the `ComBat` function in the `sva` package was used to correct for variation in the data due to the study factor [38, 39]. PCA plots were used to visualize variation in expression data before and after batch correction with ComBat [40] (Fig 2 and S1 Fig), confirming the main batch effect removal by adjusting for study.

**Fig 2.**
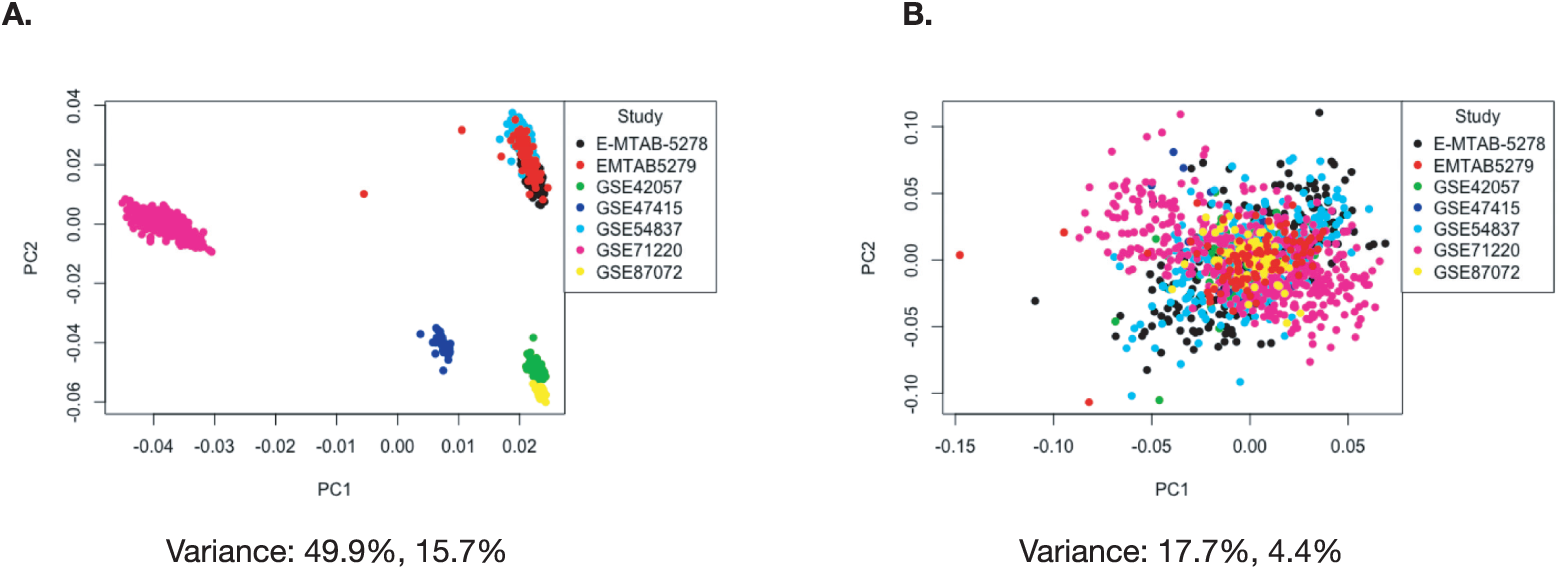
Visualizing batch effects introduced by using multiple studies in our meta-analysis. (A) PCA before and (B) PCA after batch effect correction with ComBat.

### Analysis of Variance to Identify Differentially Expressed Genes by Factor

To determine if the factors of disease status, sex, study, and smoking status had an impact on gene expression in COPD, we modeled (see linear model below) our merged expression matrix (DF2 of online supplementary data files) and then ran ANOVA to identify differentially expressed genes (Fig 1C) using `aov` and `anova` from base R’s `stats` package (as previously described [18]). Schematically our linear model formula for gene expression, *g*, per each gene included main effects and interactions:

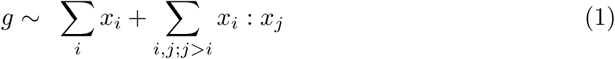

where *x*_*i*_ ∈ {age group, sex, smoker, disease status} and the factors have the following levels:

- disease status = {control, COPD}
- sex = {male, female}
- age group = {under 50, 50-55, 55-60, 60-70, over 70}
- smoker = {non-smoker, former smoker, smoker}
- study = {GSE42057, GSE47415, GSE54837, GSE71220, GSE87072, E-MTAB-5278, E-MTAB-5279}

ANOVA p-values were adjusted using the Benjamini-Hochberg (BH) correction method for multiple hypothesis testing [41–43]. Genes were considered statistically significant if their BH-adjusted p-values were <0.05. We focused on the ANOVA results for the disease factor, and filtered them for BH-adjusted p-values <0.05. These filtered genes were then identified as statistically significant disease genes. We used this gene list to identify what GO terms and Kyoto Encyclopedia of Genes and Genomes (KEGG) and Reactome pathways they were enriched in. We used the `GOAnalysis` and `KEGGAnalysis` functions from the `MathIOmica` package for GO and KEGG pathway enrichment. Additionally, we used the `enrichPathway` function from the `ReactomePA` package in R [44]. All functions for enrichment analysis used the BH p-value correction method and GO terms, KEGG and Reactome pathways with a BH-adjusted p-value <0.05 were considered statistically significant (see DF5-DF7 of online supplementary data files).

To determine the biological effect of the ANOVA statistically significant genes (disease status factor) and calculate relative expression (difference in means) to determine up- or down-regulation of genes, we conducted a post-hoc analysis with `TukeyHSD` function in the `stats` package in base R using our linear model outlined above. We added an additional column to the TukeyHSD results which contained BH-adjusted TukeyHSD p-values, and all GO terms and pathways with a BH-adjusted p-value <0.05 were considered significant. To find genes that were significantly up- and down-regulated, we further filtered the gene list by difference in means by using the two-tailed 10 and 90% quantile. With these results we carried out GO and pathway enrichment to identify which biological processes and pathways the genes were enriched. We used the disease genes and explored sex, smoking status and aging effects on their gene expression.

### Machine Learning with COPD

Machine learning classification was carried out in Mathematica using the `Classify` function [45], with the `Method` parameter set to “LogisticRegression”. We first trained on all 1262 samples, using the statistically significant disease genes, filtered with a two tailed 10 and 90% quantile selection for effect size as features (304 genes). We also randomized the dataset, and created 10 sets for training and testing, with 90% of the samples used for training, and 10% of the samples used for testing, where the 10 testing sets were mutually exclusive (10-fold cross-validation).

## Results

Our meta-analysis selection criteria for data curation (Fig 1B) resulted in 8 datasets from GEO and AE (Table 1). After pre-processing the data, we combined all datasets into a large matrix by merging by common gene names. This data merge resulted in 1,262 samples (574 controls and 688 COPD subjects) and 16,237 genes. Our 1,262 samples consists of 792 males and 470 females, and also 661 former smokers, 418 current smokers and 183 non-smokers.

### Visualizing Batch Effects and Batch Effect Correction

Prior to designing our linear model, we wanted to visualize variation introduced into the data due to batch effects, and how the variation changes when the data is adjusted with ComBat for batch effects. We used ComBat in R to adjust for the study effect on the data and generated PCA plots before and after batch correction (Fig 2). In Fig 2A, before running ComBat, the data separates into four major clusters with a variance of 49.9% in PC1 and 15.7% in PC2. After running ComBat, the clustering of the data is removed, and variance reduced to 17.7% in PC1 and 4.4% in PC2 (Fig 2B). We also plotted the PCAs for the other factors (S1 Fig) before and after using ComBat for batch effect correction. The ComBat batch effect corrected expression data was only used to visualize changes in variation due to removal of batch and to confirm the inclusion of study as an effect factor in our linear model.

### Variance in Gene Expression Due to Disease Status

With our ANOVA results, we were able to evaluate variance in gene expression introduced by each factor and their pair-wise interactions [43]. To determine which genes from our ANOVA results were statistically significant by the disease status factor, we filtered the genes by using BH-adjusted p-value <0.05. We found 1,513 statistically significant disease genes (see DF4 of online supplementary data files). We performed GO and pathway enrichment analysis on the 1,513 genes. Our enriched GO terms included: innate immune response (57 gene hits), inflammatory response (48 gene hits), apoptotic process (58 gene hits), adaptive immune response (24 gene hits) and response to drug (40 gene hits) (see DF7 of online supplementary data files for full table). We found 7 enriched KEGG pathways (Table 3 and DF5 of online supplementary data files). The enriched KEGG pathway analysis results include: Ribosome (29 gene hits), Primary immunodeficiency (11 gene hits), lysosome (22 gene hits), and cytokine-cytokine receptor interaction (35 gene hits) (Table 3 and DF5 of online supplementary data files). The 1,513 genes are involved in Reactome pathways such as Neutrophil degranulation (103 gene hits), Eukaryotic Translation Elongation (31 gene hits), Signaling by Interleukins (66 gene hits) Diseases of Immune System (8 gene hits) Fc epsilon receptor (FCERI) signaling (24 gene hits) and Signaling by the B Cell Receptor (BCR) (21 gene hits) (see DF6 of online supplementary data files for full table). We also used the `KEGGPathwayVisual` function in the `MathIOmica` package to highlight whether our gene hits for the enriched KEGG pathways were up- or down-regulated in the pathway (based on TukeyHSD calculated differences in means), Fig 3 and S2-S7 Fig. For example, Fig 3 depicts the Primary Immunodeficiency KEGG pathway and highlights our gene hits (with yellow: up-regulated, and blue: down-regulated gene expression). In this pathway, Fig 3, our results indicate that Igα is down-regulated in COPD compared to controls (involved in differentiating from a Pro-B Cell to a Pre-B cell 1), and also BTK9 is up-regulated in COPD (involved in differentiating from Pre-B1 cell to Pre-B2 cell),.

**Table 3.** Enriched KEGG Pathways using the ANOVA Differentially Expressed Genes from Disease Factor.

| KEGG ID | KEGG Pathway Name | Gene Count | *p-value* | *adjusted p-value* |
| --- | --- | --- | --- | --- |
| path:hsa03010 | Ribosome | 29 | 5.34E-08 | 1.50E-05 |
| path:hsa05340 | Primary immunodeficiency | 11 | 2.74E-05 | 3.15E-03 |
| path:hsa04142 | Lysosome | 22 | 3.37E-05 | 3.15E-03 |
| path:hsa04060 | Cytokine-cytokine receptor interaction | 35 | 1.64E-04 | 1.16E-02 |
| path:hsa04520 | Adherens junction | 14 | 4.97E-04 | 2.80E-02 |
| path:hsa05200 | Pathways in cancer | 45 | 7.00E-04 | 3.28E-02 |
| path:hsa04640 | Hematopoietic cell lineage | 15 | 9.98E-04 | 3.88E-02 |
| path:hsa05162 | Measles | 20 | 1.10E-03 | 3.88E-02 |

**Fig 3.**
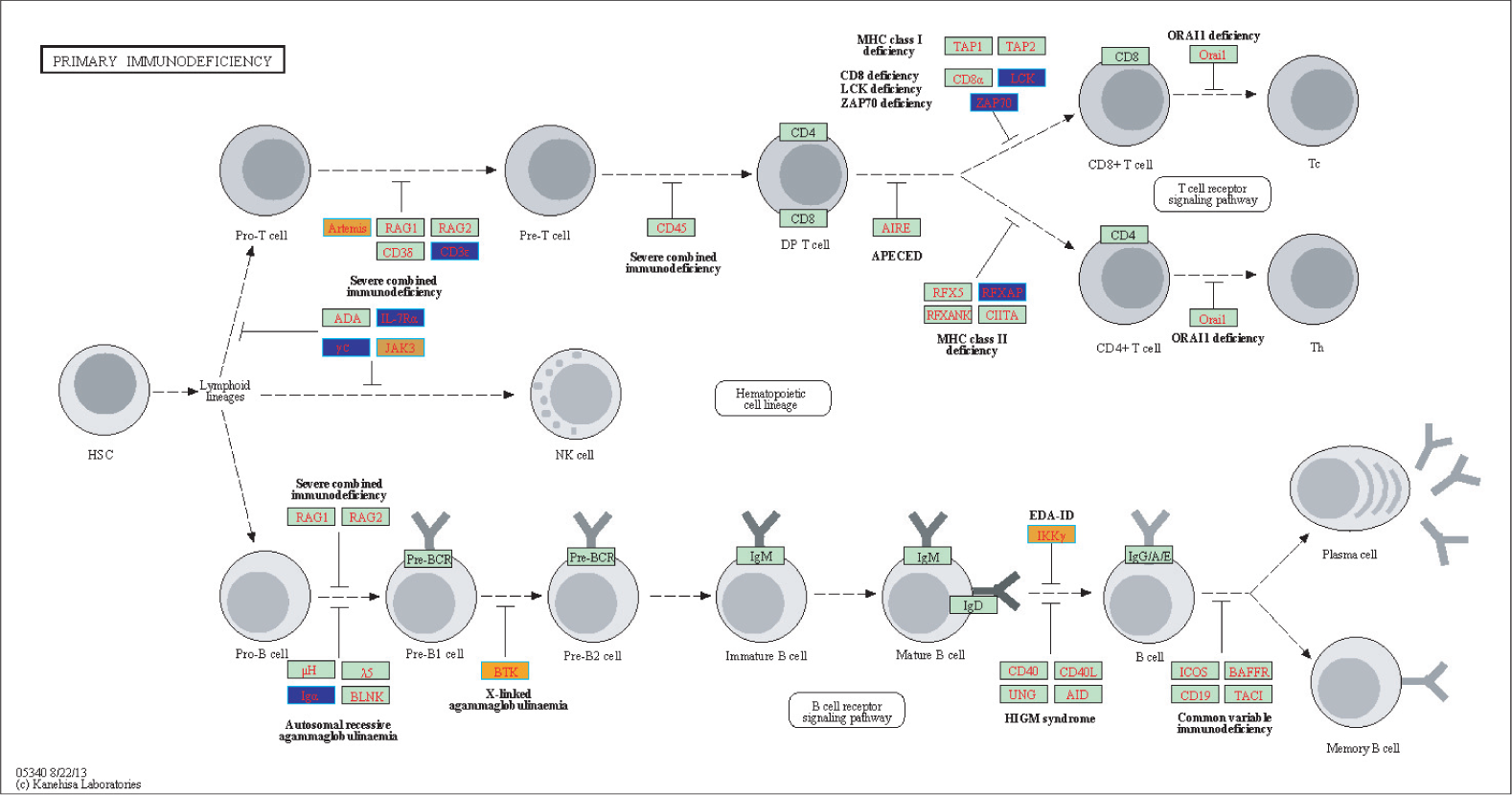
Highlighted Primary Immunodeficiency KEGG Pathway (hsa05340) with enriched genes from the ANOVA (BH-adjusted p-value < 0.05) [46–48]. Yellow-colored genes are up-regulated and blue-colored genes are down-regulated in COPD samples.

Of the 1,513 disease genes we further filtered our ANOVA results (see DF4 of online supplementary data files) to identify genes with statistically significant interactions with smoking status (disease:smoking status, BH-adjusted p-value < 0.05). We found 39 genes that had a statistically significant pairwise interaction between disease status and smoking status (see DF14 of online supplementary data files). Using the 39 interacting genes, we calculated the row means across the different pairings of smoking status and disease status to compare expression (Fig 7). We used the row means of the non-smoking controls as our baseline to calculate the difference in means for the different disease and smoking groups. In Fig 7 the data clusters by disease state (COPD together and controls together), and smokers and former smokers across both disease states have similar expression profiles. There are subset of genes that are over expressed in COPD smokers compared to control non-smokers as well as a subset of genes that are down-regulated. Finally, control smokers and former smokers have similar expression profiles with GGT6 being an outlier (Fig 7).

### Up and Down-Regulated Gene Expression in COPD

To assess biological effect and determine factorial differences in gene expression we ran TukeyHSD on our 1,513 statistically significant disease genes. We first focused on COPD and control gene expression differences and used BH-adjusted p-value <0.05 to determine significance. We also filtered further by using a 10% two-tailed quantile cutoff to identify significantly up- and down-regulated genes. Once we filtered by p-value, we calculated to 10 and 90% quantiles using differences in group means. For the COPD-control TukeyHSD comparisons we found 304 statistically significant genes that we classified as up-regulated (mean differences ⪅ −0.0260) and down-regulated (mean differences ⪆ 0.0338) in our COPD subjects. Of the 304 differentially expressed genes (DEG), 152 genes were down-regulated and 152 genes were up-regulated (DF9 of online supplementary data files). The top 25 up- and down-regulated genes are displayed in Table 4. KEGG enrichment analysis on the 152 down-regulated disease genes resulted in two significantly enriched pathways: Hematopoietic cell lineage (5 Gene hits: CD2, CD3E, CD7, FLT3LG and MS4A1) and Cytokine-cytokine receptor interaction (8 gene hits:CCL5, CCR6, CD27, CXCR3, CXCR6, FLT3LG, IL2RB, and IL2RG). For the Reactome enrichment analysis on 152 up-regulated genes, they were enriched in Neutrophil degranulation (30 gene hits) Fig 5, while the down-regulated gens were enriched in the Immunoregulatory interactions between a Lymphoid and a non-Lymphoid cell pathway (8 gene hits) (Fig 6).

**Table 4.** Top 25 up and down regulated differentially expressed genes in COPD based on effect size.

| Up-Regulated |  | Down-Regulated |  |
| --- | --- | --- | --- |
| Gene | Difference of Means | Gene | Difference of Means |
| GPR15 | 0.123 | LBH | -0.032 |
| HK3 | 0.046 | CD3E | -0.039 |
| CLEC4D | 0.075 | DUSP7 | -0.030 |
| F5 | 0.050 | TCF7 | -0.031 |
| DOCK4 | 0.063 | RRAS2 | -0.045 |
| GPR55 | 0.045 | ST6GAL1 | -0.029 |
| STAB1 | 0.043 | PYHIN1 | -0.042 |
| ASGR2 | 0.046 | CCNK | -0.026 |
| ARG1 | 0.087 | CD79A | -0.053 |
| MPO | 0.063 | GORASP2 | -0.028 |
| NRG1 | 0.064 | IL2RG | -0.029 |
| HP | 0.086 | LRIG1 | -0.028 |
| PLD1 | 0.037 | PURA | -0.027 |
| CLEC4E | 0.041 | LPAR5 | -0.033 |
| PLSCR1 | 0.048 | FAM102A | -0.030 |
| FCGR1B | 0.051 | UCP2 | -0.027 |
| FKBP5 | 0.035 | SPON1 | -0.035 |
| ANG | 0.042 | B3GNT7 | -0.042 |
| DSC2 | 0.063 | CAMK2N1 | -0.040 |
| OSBPL1A | 0.035 | DENND2D | -0.027 |
| TLR5 | 0.040 | IGFBP4 | -0.036 |
| FLVCR2 | 0.035 | IL2RB | -0.036 |
| NLRC4 | 0.037 | CD74 | -0.031 |
| AHRR | 0.063 | CBLB | -0.031 |
| CAPNS2 | 0.043 | DCXR | -0.037 |

**Fig 4.**
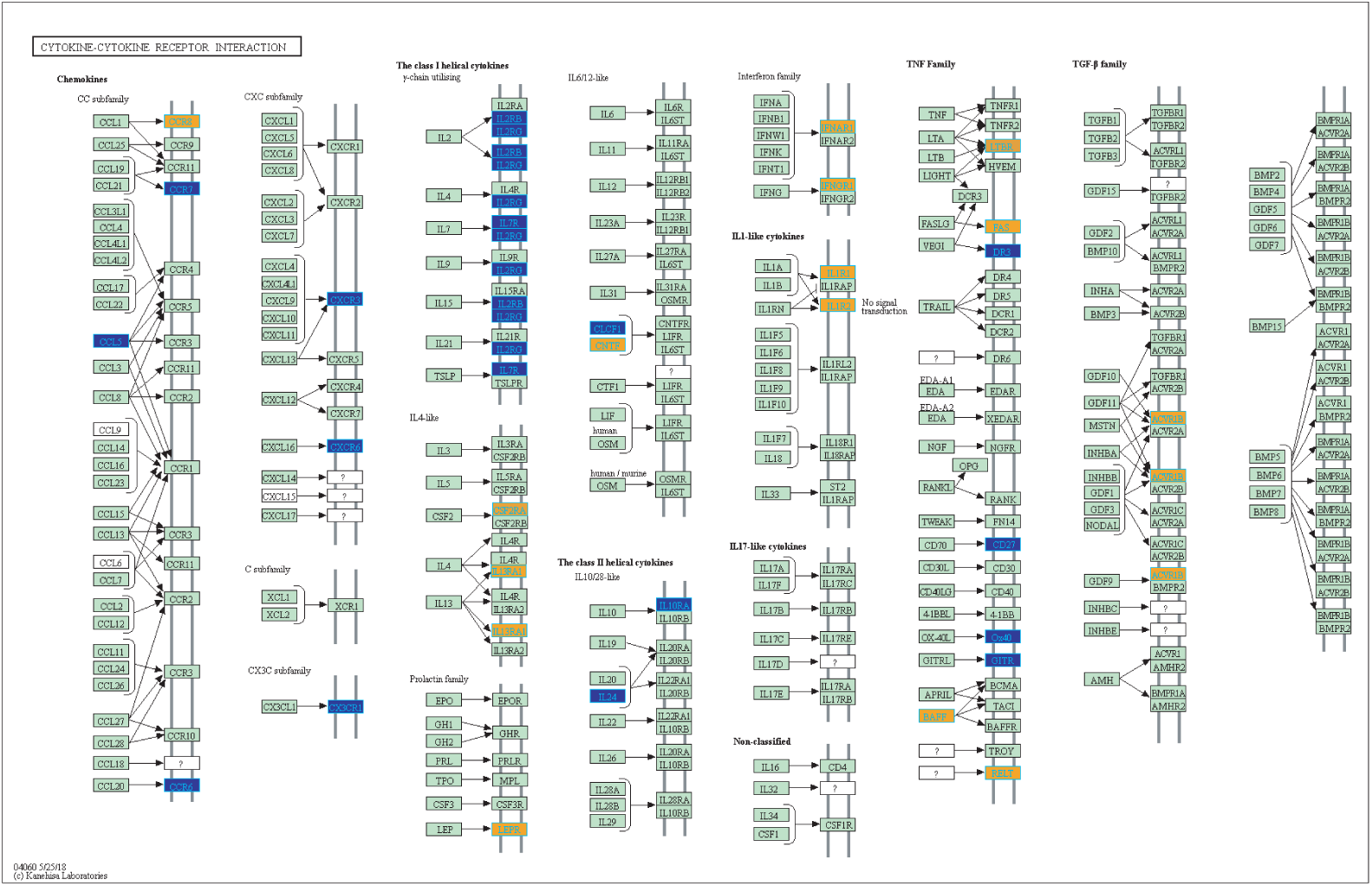
Highlighted Cytokine-cytokine receptor interaction KEGG Pathway (hsa04060) with enriched genes from the ANOVA (BH-adjusted p-value < 0.05) [46–48]. Yellow-colored genes are up-regulated and blue-colored genes are down-regulated in COPD samples.

**Fig 5.**
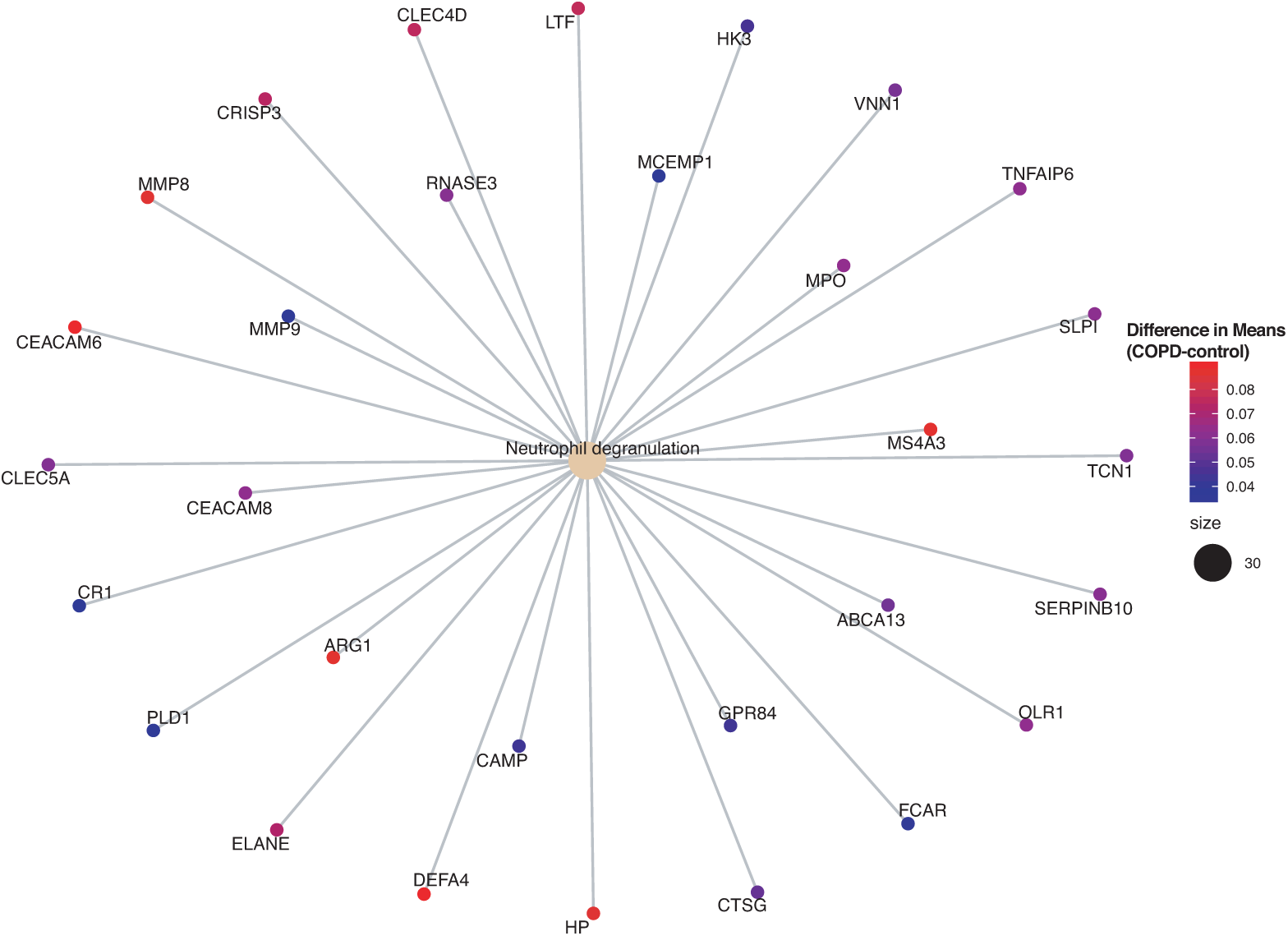
Enriched Reactome pathway-gene network from up-regulated disease genes in COPD subjects. The enrichment analysis was based on the 304 statistically significant differentially expressed genes filtered for effect size.

**Fig 6.**
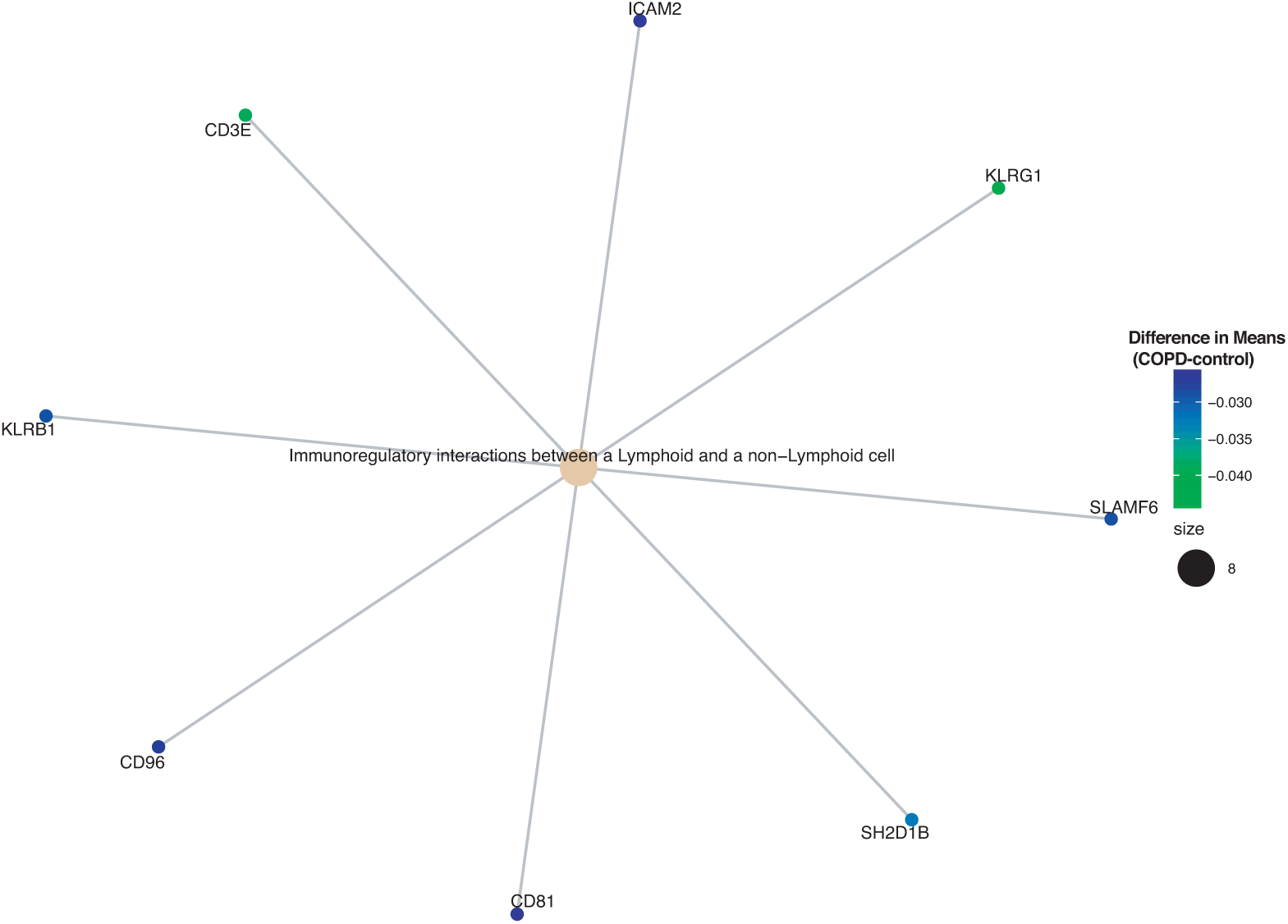
Enriched Reactome pathway-gene network from down-regulated disease genes in COPD subjects. The enrichment analysis was based on the 304 statistically significant differentially expressed genes filtered for effect size.

### Sex and Age on COPD Expression

We further analyzed the 304 DEG found to have a biological effect by disease status to identify sex and aging effects on gene expression. We found 44 genes that were differentially expressed by sex: 22 up- and 22 down-regulated in males compared to females by filtering the mean differences using two-tailed 10% quantiles, ⪅ −0.0957 (down-regulated) and ⪆ 0.0908 (up-regulated). With the 44 genes we performed pathway enrichment analysis using the `ReactomePA` package. There were 7 enriched Reactome pathways (BH-corrected p-value <0.05) that were all up-regulated in males (see DF12 of online supplementary data files and S8 Fig). These pathways include: Neutrophil degranulation (13 gene hits), Antimicrobial peptides (5 gene hits), Extracellular matrix organization (6 gene hits), Activation of matrix metalloproteinases (3 gene hits) and degradation of the extracellular matrix (4 gene hits). We also generated a gene network for the up-regulated enriched pathways in males (S9 Fig). We did not find any statistically significant interacting genes between disease status and sex from our ANOVA results.

To determine the age effect on our DEG associated with COPD (304 genes), we focused on our TukeyHSD results where the age group <50 was the baseline. We selected for significance (BH-adjusted p-value <0.05) and two-tailed 10% (up-regulated ⪆ 0.421 and down-regulated ⪅ −0.193) on the difference in means results to find significant age-group effects. We identified 304 significant age-group comparisons across 95 unique genes (see DF13 of online supplementary data files). We plotted the relative expression (difference in means) across all age comparisons with <50 as the baseline. We identified two clear clusters of the genes by expression which indicated that there are significant differences in expression profiles due to aging (Figure 8). However, we did not find any statistically significant genes with an interaction between disease status and age from our ANOVA results.

**Fig 7.**
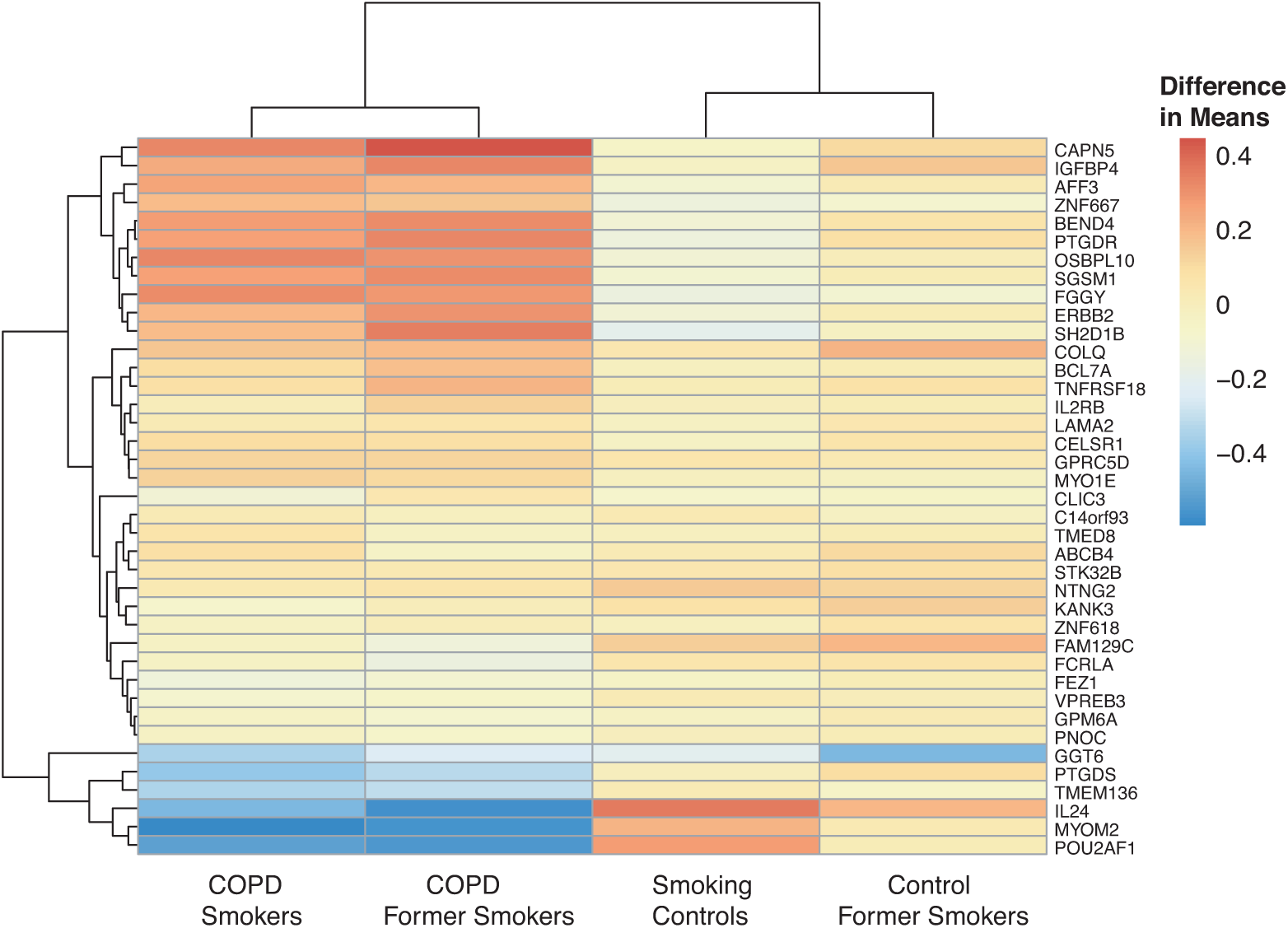
Heatmap of statistically significant interacting genes across disease states and smoking statuses. Difference in means calculated using control non-smokers as the baseline.

**Fig 8.**
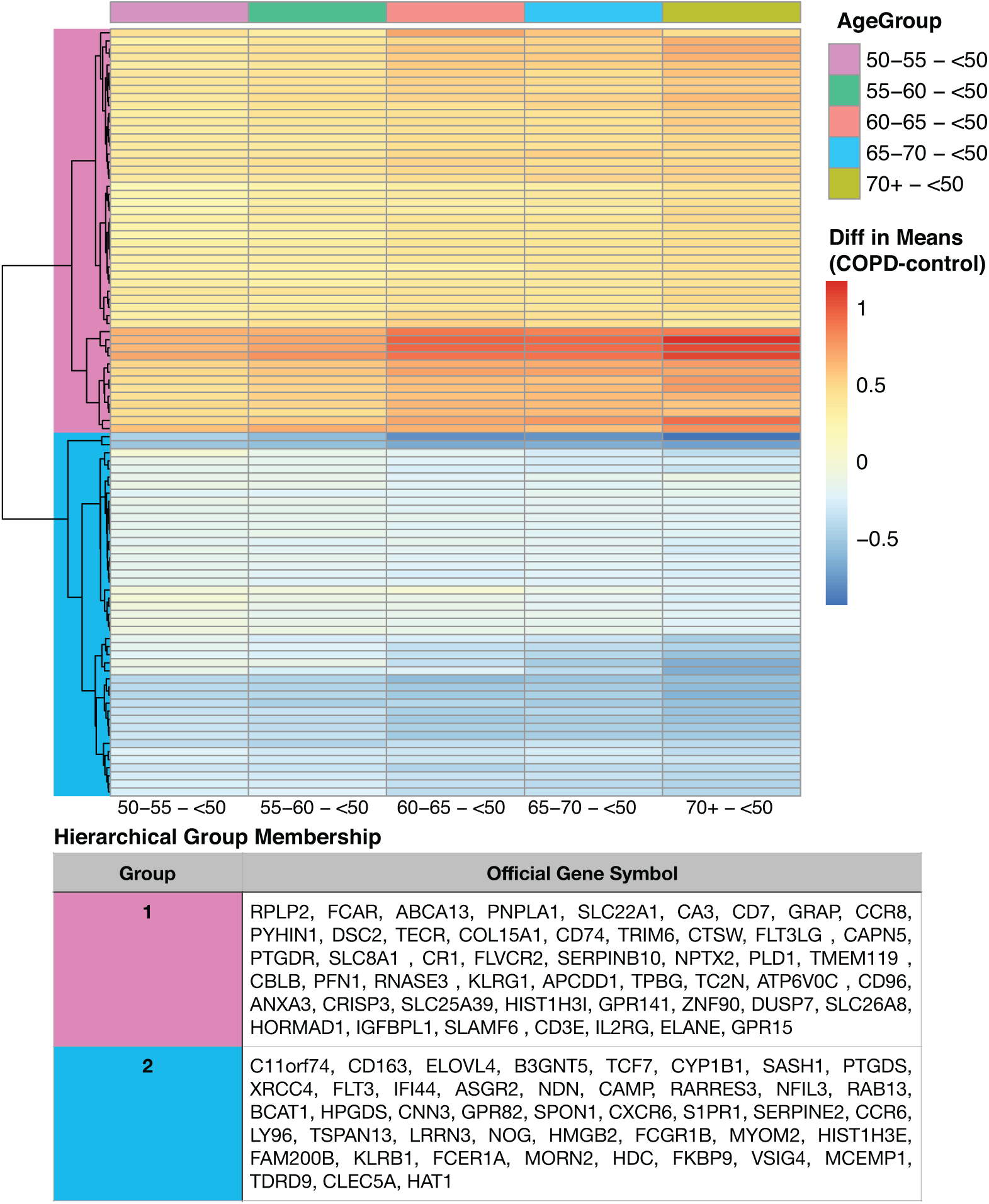
Heatmap of age effect on the statistically significant disease gene list. The enrichment analysis was based on the 304 statistically significant differentially expressed genes filtered for effect size. The clustered groups are color-coded, with the corresponding genes in each group listed in the table.

### Machine Learning with COPD Data

Using the gene expression from the top 304 statistically significant for disease genes, and with 10% two-tailed highest effect size we trained a logistic regression model in Mathematica for predicting whether a profile belongs to the control or COPD group. Training with all samples achieved an accuracy of 87.0 *±* 3.0%,(Fig 9A). The corresponding confusion matrix and receiver operating characteristic (ROC) curves are shown in Fig 9 respectively, with an ROC area under the curve (AUC) of 0.979. Furthermore, we decided to carry out a 10-fold cross-validation analysis of randomized order samples, where we trained on 90% of the data each time and tested on the remaining 10%. On average the model had an accuracy of 84.2% (standard deviation of 3.1%), and ROC AUC of 0.921 (standard deviation of 0.022). An example of the worst performing realization from the cross-validation is shown in Fig. 9D-F, where 48/57 controls and 42/69 COPD samples were classified correctly, whereas 9/57 controls were mis-classified as COPD, and 17/69 COPD were misclassified as controls. Equivalently, the false positive rates were on average 0.17 (control) and 0.14 (COPD), and the false discovery rates were on average 0.19 (control) and 0.12 (COPD).

**Fig 9.**
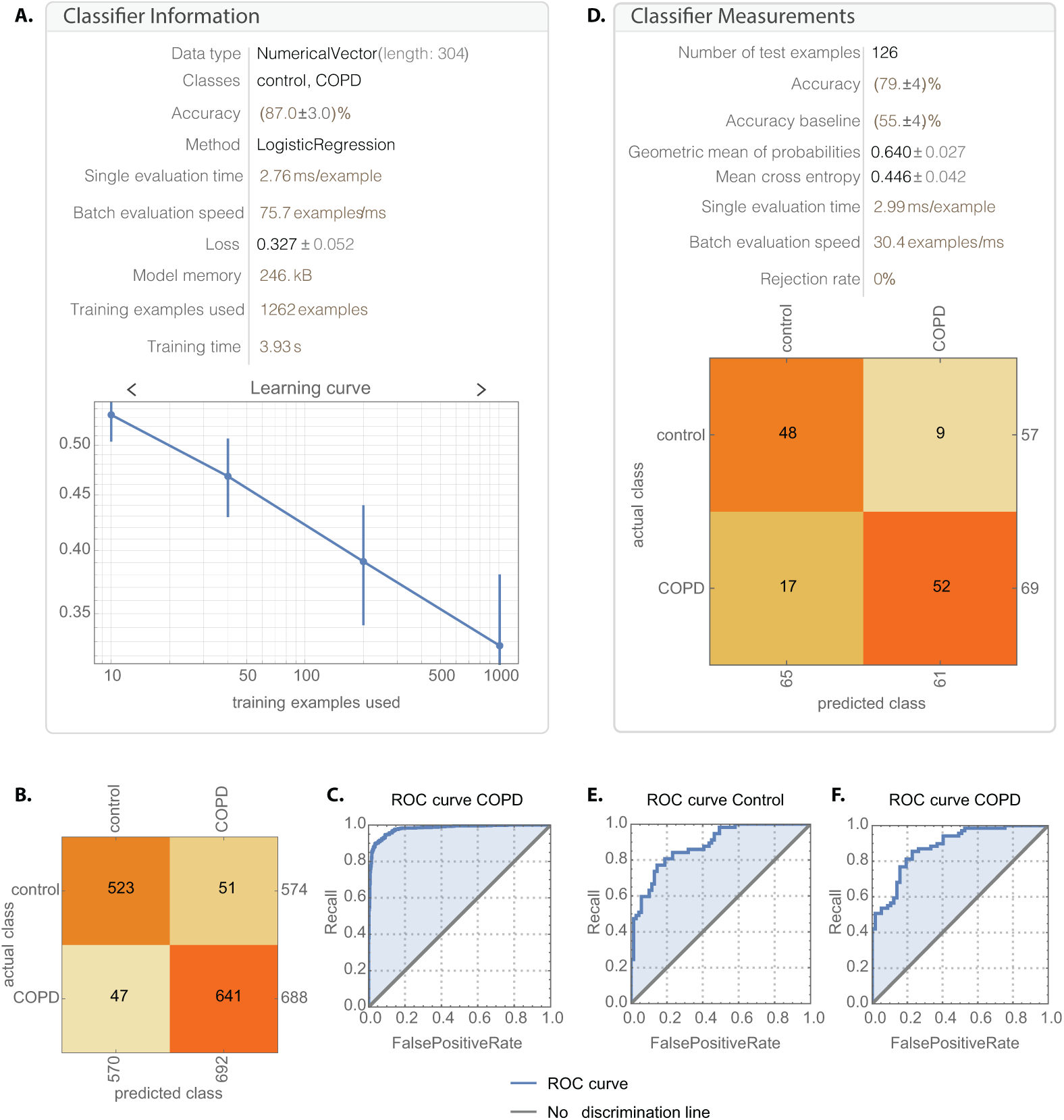
Trained logistic regression model can classify COPD and healthy profiles. (A)The logistic regression model trained on all the data achieves 87.0 *±* 3.0% accuracy), with the (B) confusion matrix and (C) ROC curves indicating good performance overall, with AUC 0.979. Training with 10-fold cross validation gives an average accuracy of 84.2%, with the worst testing model shown in (D) and its ROC for Controls and (F) COPD shown respectively, with an AUC of 0.882.

## Discussion

Chronic obstructive pulmonary disease causes damage to the lungs because of exposure to toxic irritants or genetic factors, and is a rising global health problem. With an increase in the elderly population’s life expectancy and the number of smokers, the prevalence of COPD and its morbidity rates are expected to rise. Researchers are working to identify strategies that can help to clearly understand COPD, its pathology, and to find biomarkers in easily accessible body fluids to promote earlier detection of COPD and improve accuracy of diagnosis [15–17]. Our research objective was to identify age, sex and smoking status effects on gene expression between COPD and controls in blood. We curated and downloaded 7 microarray expression datasets for our meta-analysis on COPD. Using the raw expression data, we removed the background, annotated and summarized the probes, and merged the 7 datasets together by common gene names. This was followed by data normalization using BoxCox power transformation and downstream analyses to identify differentially expressed genes and genes that were biologically significant. This is the largest COPD meta-analysis and explores expression variability in 1,262 samples by modeling linear and binary effects of disease status, age, sex and smoking status.

Our ANOVA highlighted 1,513 statistically significant (BH-adjusted p-value <0.05; disease status factor) disease genes (see DF4 of online supplementary data files). One of our genes, FAM13A, has previously been associated with COPD susceptibility [8, 49]. Other genes such as GPR15, CLEC4D and MPO have also been associated with COPD and inflammation within the lungs. Our GO and pathway enrichment results highlight some immune pathways (Table 3) and GO terms such as innate immune response, adaptive immune response and inflammation (DF7 of online supplementary data files) that have previously been associated with COPD. For example, primary immunodeficiency (weakened immune system due to deficiencies in immune cell production) is linked to recurrent infections in subjects with COPD [50]. This recurrence in infections due to a weakened immune system also causes chronic inflammation and airway remodelling and obstruction [50]. Humoral deficiencies and inadequate antibody production and responses to infections also reduce the effectiveness of vaccinations such as the influenza and pneumococcal vaccines [50, 51]. Studies have suggested that antibody replacement therapy can help to reduce the recurrence of bacterial and viral infections in COPD subjects [50, 52]. In our results, we highlighted the primary immunodeficiency KEGG pathway to determine how our genes are regulated in the pathway (Fig 3). Of the 8 genes highlighted in the T cell maturation portion of Fig 3, 6 of them are down-regulated in COPD subjects compared to our controls. IL2RG (alias gamma chain(γC)) that regulates T cell development and differentiation was down-regulated in COPD subjects (Fig 3). IL2RG is also associated with severe combined immunodeficiency [53]. Previous findings suggest that down-regulation of the soluble common gamma chain is a mechanism to reduce inflammation by T cells in response to cigarette smoke in a COPD mouse model [54]. Up-regulated γC promotes interferon-γ production and inflammation in the respiratory tract [54]. IL7-Ra and JAK3 are also linked to severe combined immunodeficiency. DCLRE1C (Artemis) and CD3E are both involved in Pro-T to Pre-T cell differentiation and were up- and down-regulated respectively in COPD subjects (Fig 3). Genes such as LCK ZAP70 and RFXAP are involved in T cell differentiation into CD8+ and CD4+ cells and were found to be down-regulated in COPD (Fig 3). In B-cell differentiation, our gene hits, BTK (B-cell development) and IKBKG (alias IKKγ) were up-regulated in COPD while Igα was down-regulated (Fig 3). Reduced Igα or deficiencies in Igα promote reoccurring infections and disease exacerbation in COPD subjects [50, 55].

In the highlighted Cytokine-cytokine receptor interaction KEGG pathway there are different classes of cytokines such as chemokines, class I cytokines and the Tumor necrosis factor and Transforming growth factor beta families with varying expression (Fig 4). Cytokines play a major role in the inflammatory response observed in COPD subjects. For instance, CCR8 (chemokine) was up-regulated in COPD subjects (Fig 4). Increased levels of CCR8 has been previously observed in allergic asthmatics [56] and has a functional role in macrophage processes and release of cytokines in the lungs [57].

We also visualized our up- and down-regulated gene hits in the other enriched KEGG pathways (Table 3 and S2 - S6 Fig). We highlighted our 45 gene hits in the Pathways in Cancer KEGG pathway (S2 Fig). COPD is a known risk factor for lung cancer and it leads to 1% of cancer cases each year [58]. Furthermore, there is a five-fold increase to developing lung cancer in patients with COPD compared to individuals with normal pulmonary function [58]. Some of our highlighted genes are involved in apoptosis (Fas and CASP9), DNA damage (MDM2), Extra-cellular matrix (ECM) receptor interaction (ECM) and proliferation (CyclinD1) (S4 Fig). As for the KEGG Lysosome pathway (S3 Fig), lysosome function and distribution in the cells of COPD subjects and smokers have been previously examined. The lysosomes in smokers have been previously shown to cluster around the nucleus of the cell and with reduced concentrations of lysosomes throughout the cell compared to subjects who did not smoke. Additionally, dysregulation of the lysosomal pathway has also been previously described in COPD patients [59].

We observed some down-regulated genes in the adherens junction pathway for COPD subjects (S4 Fig). This may be connected to the increase in lung epithelial permeability due to smoking. Also, one study highlighted that apical junctional complex (AJC) genes were down-regulated in COPD smokers, and that the cigarette smoke promotes a cancer-like molecular phenotype by causing reprogramming of transcription of the AJC [60]. The hematopoietic cell lineage pathway highlights genes involved in the differentiation of immune cells from hematopoietic stem cells (S5 Fig). As for the enriched measles pathway, research suggests that heavy smokers who had childhood measles has an increased risk for developing COPD [61]. The Reactome pathway analysis also resulted in immune related pathways such as Neutrophil degranulation, Signaling by Interleukins, Diseases of the Immune System and Signaling by the B Cell Receptor which all highlight components of the pathology of COPD (DF6 of online supplementary data files).

Focusing on the 304 differentially expressed disease genes (filtered for biological effect), some of the top up-regulated genes are GPR15 (found on lymphocytes and involved in trafficking of lymphocytes), HK3 (glucose metabolism), CLEC4D (role in inflammation and immunity) and F5 (blood coagulation factor) [53] (Table 4. As for our top down-regulated genes CD3E (role in T-cell development), DUSP7 (involved in MAPK signaling), TCF7 (role in natural killer cell development), RRAS2 (involved in cell proliferation). We also wanted to compare our gene list to a previously published meta-analysis. Reinhold et al., had a total of 6,243 genes which they grouped into 15 modules for each cohort [17]. Out of our 304 genes, 97 of them overlapped with their findings while 207 of our genes were unique. We used BINGO in Cytoscape v.3.7.1 for GO analysis on our 207 unique genes (Fig S9 and S3 File) [62, 63]. Our BINGO results (BH-adjusted p-value < 0.05) include GO terms such as defense response, response to bacterium, response to stress, response to wounding, immune response, cell adhesion, and inflammatory response (Fig S9 and S3 File).

In addition to exploring enriched GO terms associated with our 304 disease genes, Fig 5 - 6 highlight the genes that were up-regulated in COPD and were enriched in the Reactome pathways Neutrophil degranulation (Fig 5) (genes up-regulated in COPD), and Immunoregulatory interactions between a Lymphoid and a non*–*Lymphoid cell pathway (genes up-regulated in COPD). Neutrophil degranulation (release of granules by exocytosis) has been associated with pulmonary disorders including asthma and COPD. In COPD patients’ neutrophils are the highest number of inflammatory cells present in the bronchial walls [64]. Increase neutrophil degranulation induces tissue damage and this is due to high inflammatory state and constant priming of neutrophils by cytokines and chemokines [64]. Our up-regulated genes in the neutrophil granulation pathway include CEACAM6 (cell adhesion), MMP8 (tissue remodeling and breakdown of extracellular matrix), CLEC4D (cell-adhesion, cell signaling and inflammation), LTF (granules in neutrophils), MS4A3 (signal transduction), and DEFA4 (defense antimicrobial peptides). Immunoregulatory interactions between a Lymphoid and a non*–*Lymphoid cell pathway down-regulated genes include KLRB1 and KLRG1 (role in the regulation of natural killer cell function), CD3E (involved in adaptive immune response), ICAM2 (leukocyte adhesion and recirculation), SLAMF6 (natural killer cell activation) and CD81 and CD96 (role in adaptive immunity) [53].

To assess the effect of smoking status on gene expression, we focused on the biologically significant genes with a significant interaction between disease status and smoking status. We identified 39 disease genes that significantly interacted with smoking status (Fig 7). The baseline in Fig 7 was non-smoking controls. For the two control groups: current and former smokers, they both have elevated gene expression levels compared to non-smoking controls. This indicates changes due solely to smoking with moderate differences between former and current smokers. As for the COPD smokers and non-smokers, the majority of these genes are elevated compared to non-smoking controls with GGT6, PTGDS, TMEM136, IL24, MYOM2 and POU2AF1 being down-regulated in COPD compared to healthy non-smokers. Some of these genes have been associated with lung function and disorders such as GCT6 which plays a role in gluthathione homeostasis and lung airspace epithelial barrier [65], IL-24 can induce apoptosis and helps control cancer cells [66] and POU2AF1 is a regulator of host defenses but cigarette smoke suppresses its gene expression [67] (Fig 7). In our analysis there was only 1 COPD non-smoker which was excluded from this analysis.

As for sex specific effects on gene expression, we identified 44 of the 304 disease genes to have a sex effect. The enriched pathways from using the genes that were up-regulated in males are highlighted in S7 Fig. These genes are involved in Reactome pathways such as Neutrophil degranulation, Extracellular matrix organization, Collagen degradation, Degradation of the extracellular matrix, and antimicrobial peptides (S7 Fig). Neutrophil degranulation was discussed above as being up-regulated by disease status in COPD subjects compared to controls. In COPD, the extracellular matrix of the airway and parenchyma of the lungs are restructured [68, 69]. Previous findings observed altered expression of elastin and collagen in COPD compared to controls, and the stage/severity of COPD affected extracellular matrix remodeling [68, 69]. Studies on COPD and sex, previously suggested higher prevalence in males due to them having higher smoking rates [70, 71]. However, currently with larger numbers of women smoking the prevalence of COPD in women is on the rise. Studies have shown that women are 50% more susceptible to COPD than males and why this is the case is still an on going debate [70, 71]. Some reasons include, smaller airways so larger concentrations of tobacco smoke in the lungs and hormonal effects [70, 71]. Of the 44 genes with a sex effect, we did not find any genes with a significant interaction between disease status and sex.

Aging trends were visualized on the biologically significant disease genes. 95 genes showed significant aging trends compared to our baseline (<50) (DF13 of online supplementary data files). Symptoms for COPD can be detected between ages 40 and 50 [72], and because of this we used our subjects grouped as <50 as our baseline. The data clustered into two distinct groups with similar gene expression patterns (Fig 8). Group 1 genes were significantly up-regulated for all age groups compared to the baseline (Fig 8). Pathway enrichment analysis indicated that the genes in group 1 are involved in the Neutrophil degranulation pathway (p-value 1.06e-04 and FDR 0.029) which has previously been described above as being up-regulated in COPD subjects. Genes within group 2 displayed an opposite trend with most genes being down-regulated with increasing age (Fig 8). These genes did not result in any statistically significant enrichment. However, genes in group 2 include C11orf74 (involved in transcription regulation) [53], CD163 (previously found to be over expressed in lungs of individuals with severe COPD) [73], TCF7 (natural killer and lymphoid cell development) [53], CYP1B1 (previously shown to be up-regulated in COPD and smokers) [74] and SASH1 (involved in TLR4 signaling and can promote cytokine production) [53]. In addition to this, we did not find any significant interacting genes between disease status and age.

To test the possibility of using blood expression data from micro-arrays to predict disease status, we performed machine learning with a logistic regression model using the 304 disease genes. This resulted in an average accuracy of 84.2% (Fig 9). These results are promising despite using aggregate expression versus cell-type specific expression. Previous studies explored using computed tomography (CT) images COPD patients and controls for disease classification [75]. Some studies also used patient reported data (such as heart rate, respiratory rate) to predict disease exacerbation and resulted in an ROC of 0.87 [76] and another with 70% sensitivity and 71% specificity [77].

Conducting a meta-analysis with microarray expression data limits our findings to annotated genes, and hinders us from discovering novel genes and looking at the entire transcriptome. Additionally, using publicly available data limits us to specific factors we can explore in our analysis due to subject characteristics not being reported uniformly across datasets (see S1 File). For example, all studies did not report ethnicity and therefore we could not investigate the effect of ethnicity on gene expression in COPD. This would be a good factor to explore due to over 90% of COPD cases occurring in low-middle class communities [5, 10]. We also did not have consistently reported disease severity information to factor into our analysis and findings. Our selection criteria for the publicly available data limits our sample size (Fig 1). In addition to this, the limitations of available data resulted in unbalance in sample constitution: 1,262 samples with 574 controls and 688 COPD, of which 792 are males and 470 females, and have smoking status as 183 non-smokers, 418 smokers, and 661 former smokers. As for our machine learning algorithm, despite having a good predictive power and accuracy, we could not explore cell-type specific data. Furthermore, the observed confounding between studies suggests that samples would need to be analysed together with the current sample sets in new investigations, prior to prediction of status.

Our study highlights new gene candidates by factor (disease status, age, sex and smoking status) and genes that statistically interact between disease status and smoking status that can be studied further to understand their role in COPD. Future work to expand on our findings must include the use of cell-type specific expression data and RNA-sequencing data. Due to COPD being characterized by inflammation, increased macrophages and neutrophils and their release of cytokines, looking at cell-type specific data can give more insight on pathology of COPD. Using cell-type specific data for predicting disease states will also expand on our findings. RNA-sequencing data can introduce novel gene candidates and biomarkers for COPD. Furthermore, implementing proteomics and metabolomics can help characterize disease pathology and may lead to discovery of additional signatures for early detection of COPD using a systems biology approach.

## Data Availability

Online Supplementary Data files. All of our datasets, data files and results from our COPD meta-analysis have been deposited to FigShare. The file names begin with the prefix “DF” and are referred to throughout the manuscript. To access our supplemental data files access the FigShare online repository at: https://doi.org/10.6084/m9.figshare.8233175. Datasets used in the meta-analysis are available from Gene Expression Omnibus and Array Express, and their accessions are listed in Table 1.

## Supporting information

**S1 Fig. Principal Component Analysis to visualize changes in variation in datasets before and after combat**. (A) PCA using disease state as batch before ComBat, (B) PCA on disease state after ComBat, (C) PCA using sex as batch before ComBat (D) PCA on sex after ComBat (E) PCA using smoking status as batch (F) PCA on smoking status after ComBat.

**S2 Fig. Highlighted Pathways in Cancer KEGG Pathway with enriched genes from the ANOVA (BH-adjusted p-value** < **0.05; disease status factor) [46–48]**. This figure highlights the relative expression (difference in means) of the gene hits from our data on the Pathways in Cancer pathway. Genes colored in yellow are up-regulated and blue-colored genes are down-regulated in COPD. The enriched genes are from the ANOVA statistically significant disease status gene list (BH-adjusted p-value < 0.05, disease status factor). Our TukeyHSD (used to determine up- or down-regulation of genes) results are with our online data files (DF16 of online supplementary data files).

**S3 Fig. Highlighted Lysosome KEGG Pathway with enriched genes from the ANOVA (BH-adjusted p-value** < **0.05; disease status factor)**. This figure highlights the relative expression (difference in means) of the gene hits from our data on the Lysosome pathway. Genes colored in yellow are up-regulated and blue-colored genes are down-regulated in COPD. Our TukeyHSD (used to determine up- or down-regulation of genes) results are with our online data files (DF17 of online supplementary data files).

**S4 Fig. Highlighted Adherens KEGG Pathway with enriched genes from the ANOVA (BH-adjusted p-value** < **0.05; disease status factor) [46–48]**. This figure highlights the relative expression (difference in means) of the gene hits from our data on the Adherens pathway. Genes colored in yellow are up-regulated and blue-colored genes are down-regulated in COPD. Our TukeyHSD (used to determine up- or down-regulation of genes) results are with our online data files (DF18 of online supplementary data files).

**S5 Fig. Highlighted Hematopoietic Cell Lineage KEGG pathway with enriched genes from the ANOVA (BH-adjusted p-value** < **0.05; disease status factor) [46–48]**. This figure highlights the relative expression (difference in means) of the gene hits from our data on the Hematopoietic Cell Lineage pathway Genes colored in yellow are up-regulated and blue-colored genes are down-regulated in COPD. Our TukeyHSD (used to determine up- or down-regulation of genes) results are with our online data files (DF20 of online supplementary data files).

**S6 Fig. Highlighted Measles KEGG pathway with enriched genes from the ANOVA (BH-adjusted p-value** < **0.05; disease status factor) [46–48]**. This figure highlights the relative expression (difference in means) of the gene hits from our data on the Measles pathway. Genes colored in yellow are up-regulated and blue-colored genes are down-regulated in COPD. Our TukeyHSD (used to determine up- or down-regulation of genes) results are with our online data files (DF21 of online supplementary data files).

**S7 Fig. Enriched Reactome pathway-gene network using the differentially expressed disease genes with a sex effect (no significant interaction between sex and disease) that were up-regulated in males)**. The gene list used and the pathway results can be found in DF11-12 of online supplementary data files.

**S8 Fig. Gene ontology results from BINGO using our 207 unique statistically significant disease genes filtered for biological effect**. Our 304 biologically significant genes were compared to Reinhold et al., [17] We ran GO with BINGO on 207 of our unique genes. The node size relates to number of genes, and the yellow nodes are statistically significant with a BH-adjusted p-value <0.05 and FDR <0.05.

**S1 File. Datasets and the information reported on samples used for the meta-analysis**. This Microsoft Excel file lists all of the studies included in the meta-analysis ans well as their sample description and study details. It highlights factors not commonly reported across all datasets.

**S2 File. Description of our online supplementary data**. This Microsoft Excel file lists all of our supplemental data files (datasets and results) from our meta-analysis. See our data availability statement for more information.

**S3 File. Gene ontology results from BINGO**. Table of the GO results from our 207 unique disease genes from our comparison of results with the findings of Reinhold et al. [17].

