## Supplementary figures and images for "Meta-analysis of Gene Expression Microarray Datasets in Chronic Obstructive Pulmonary Disease"

### S1 Fig

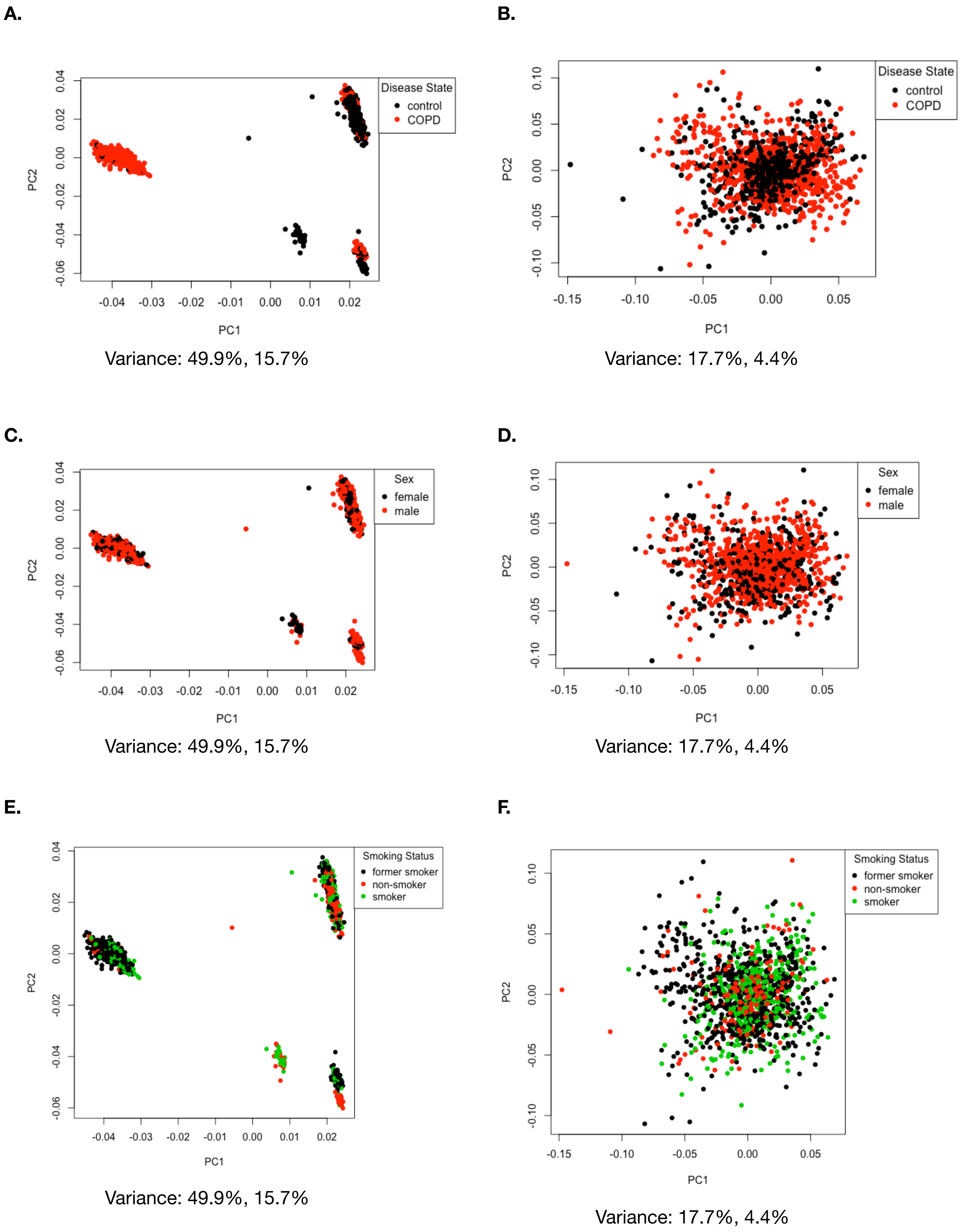

### S2 Fig

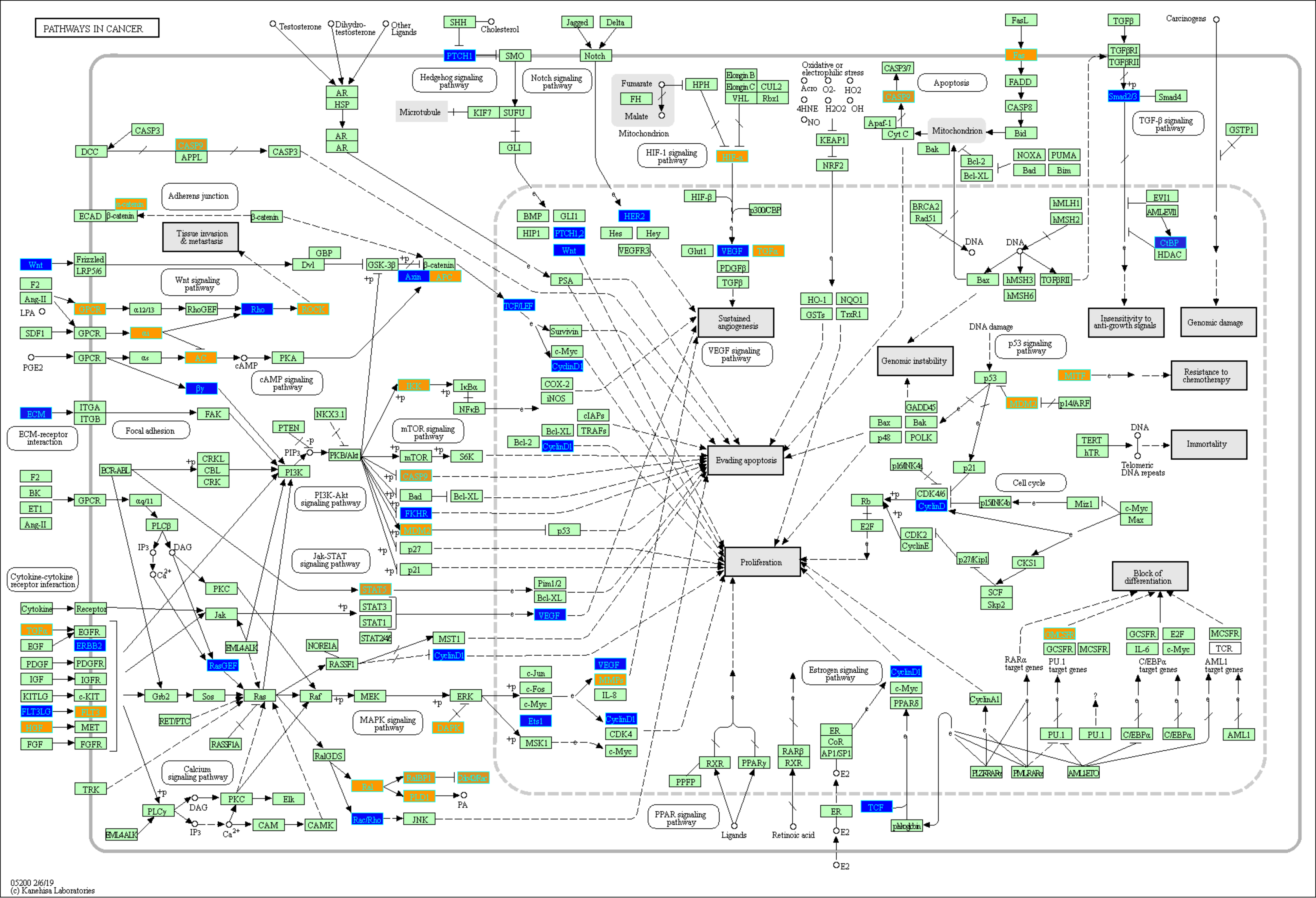

### S3 Fig

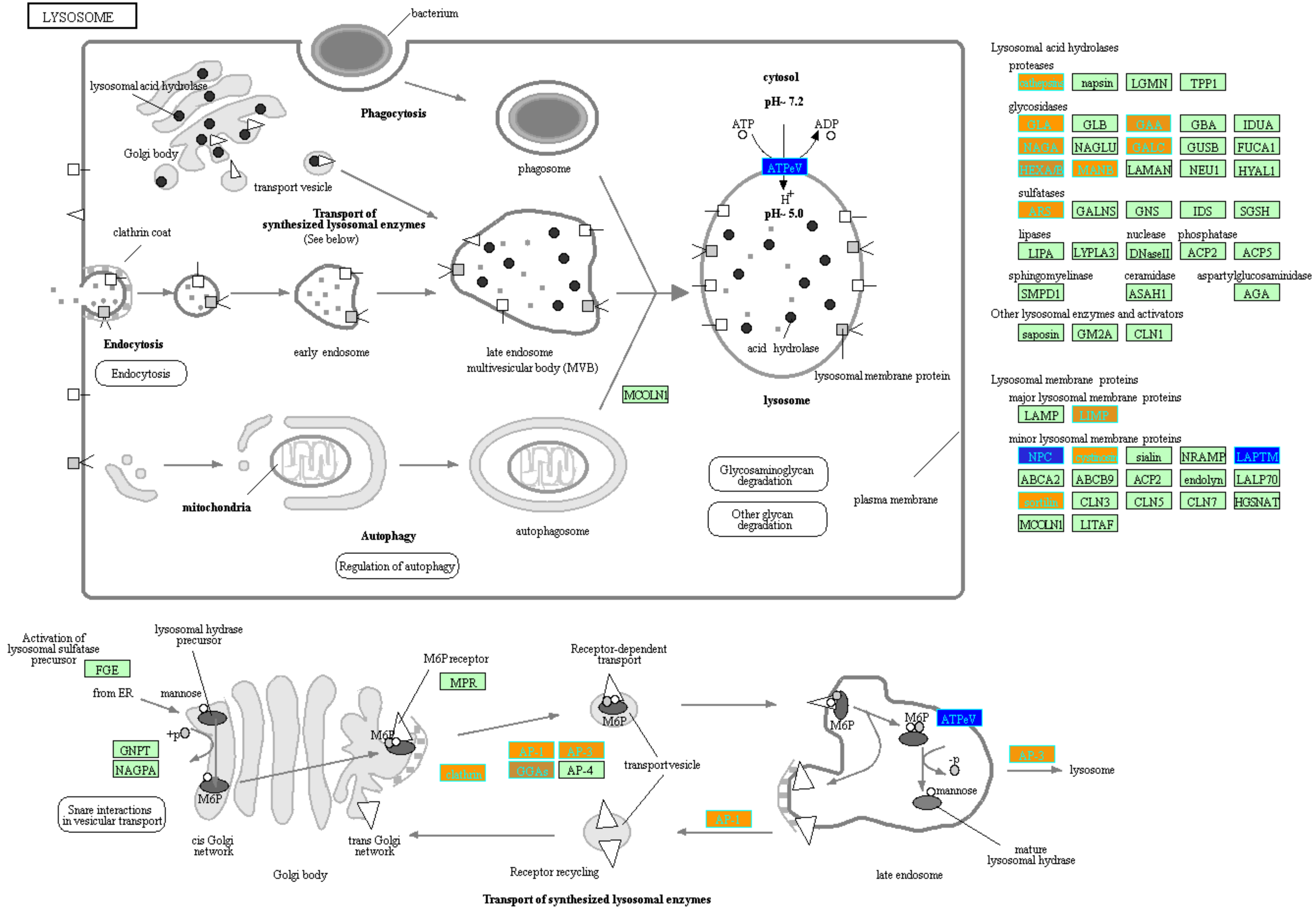


04142 3/16/17
(c) Kanehisa Laboratories

### S4 Fig

ADHERENS JUNCTION

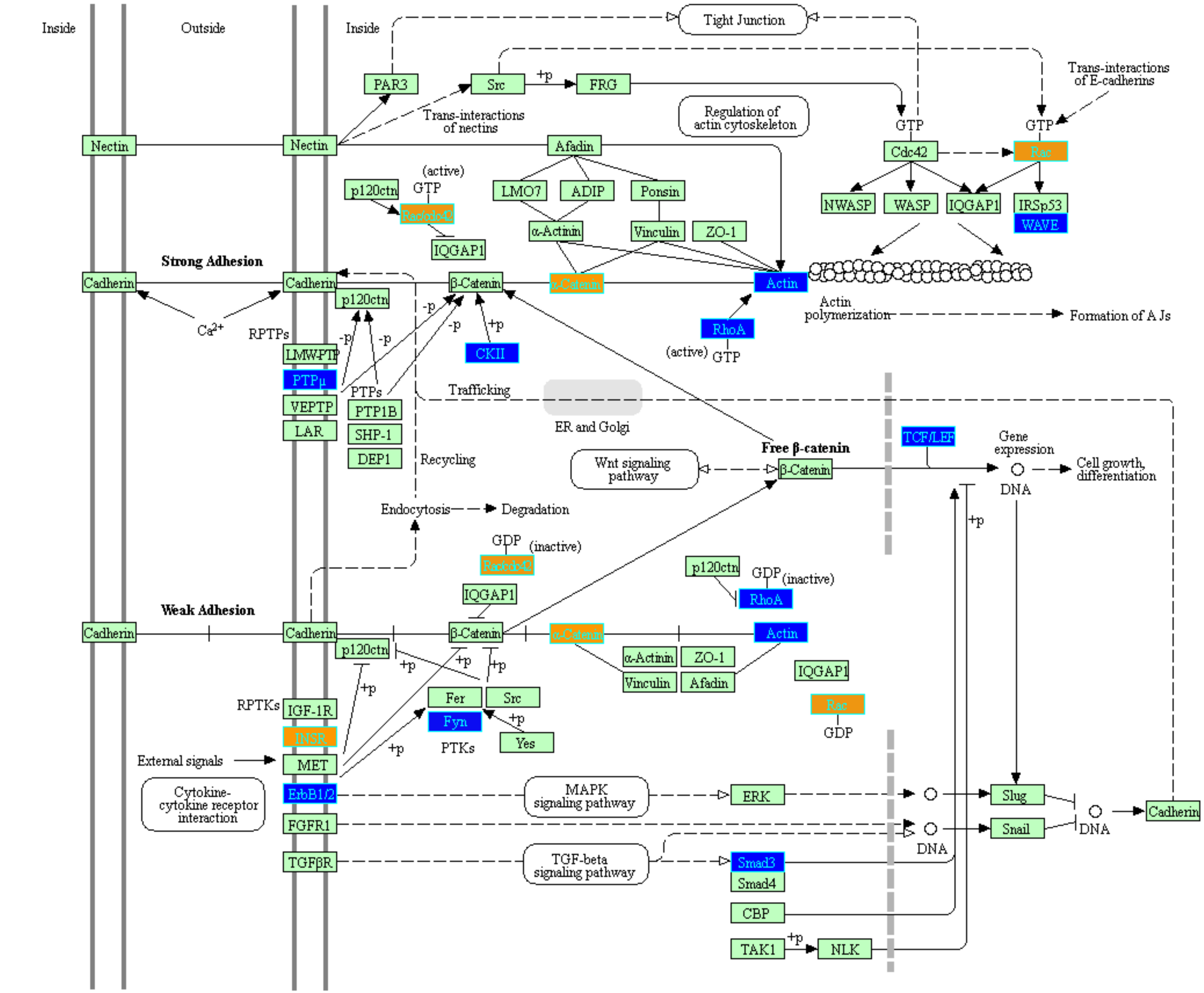


04520 3/1/13

(c) Kanehisa Laboratories

### S5 Fig

#### HEMATOPOIETIC CELL LINEAGE

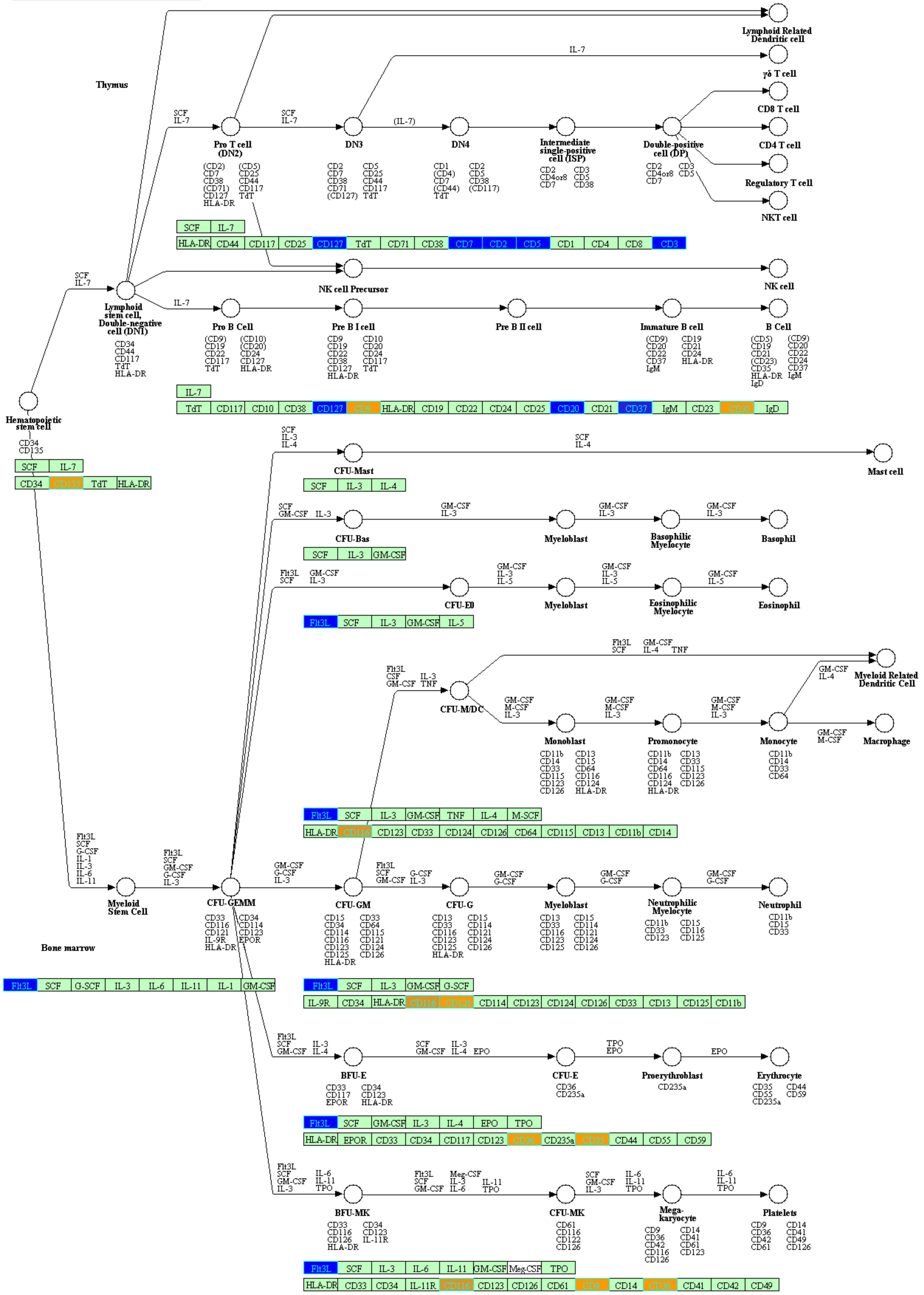


04640 10/31/16
(c) Kanehisa Laboratories

### S6 Fig

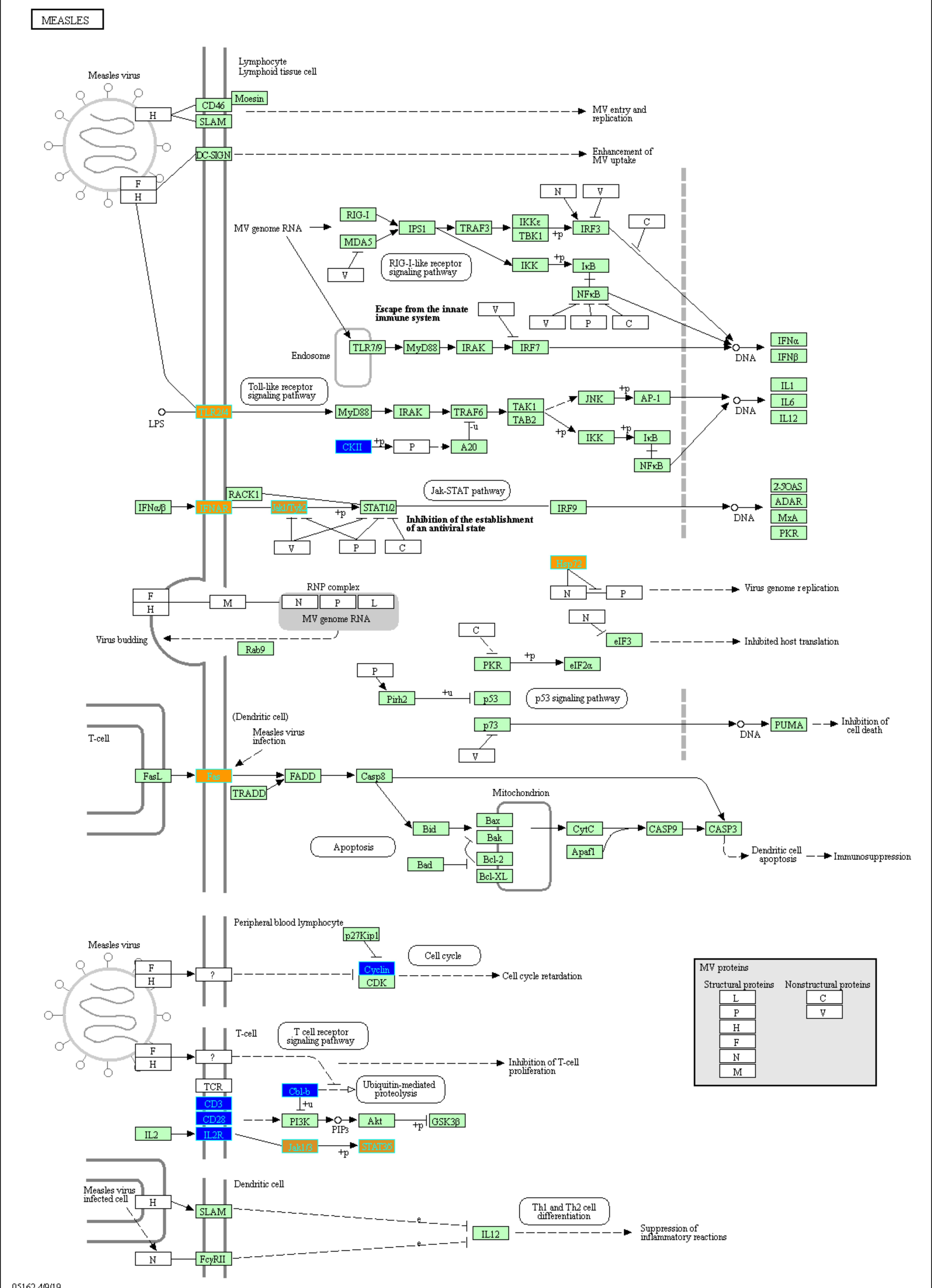


05162 4/9/19
(c) Kanehisa Laboratories

### S7 Fig

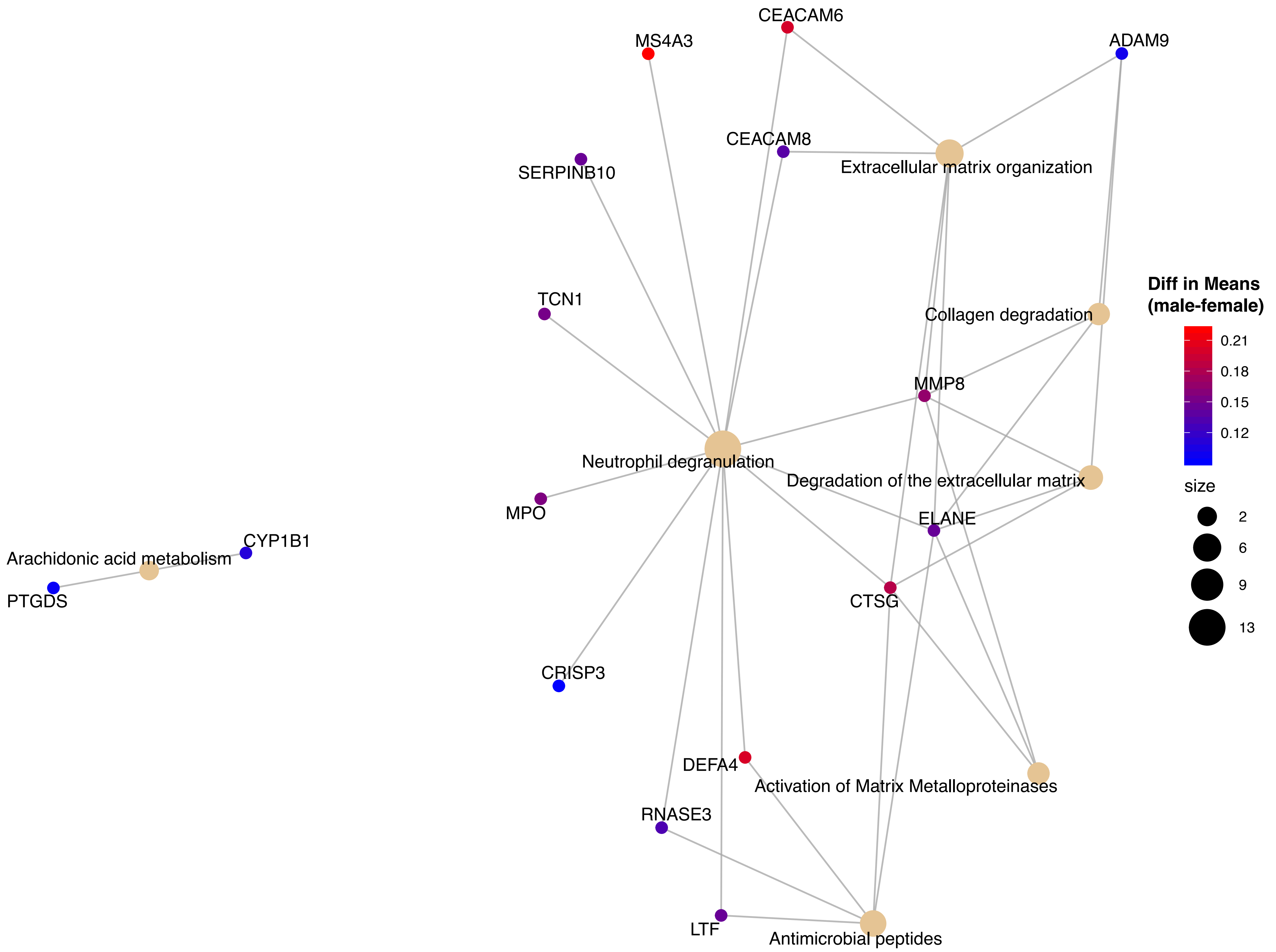

### S8 Fig

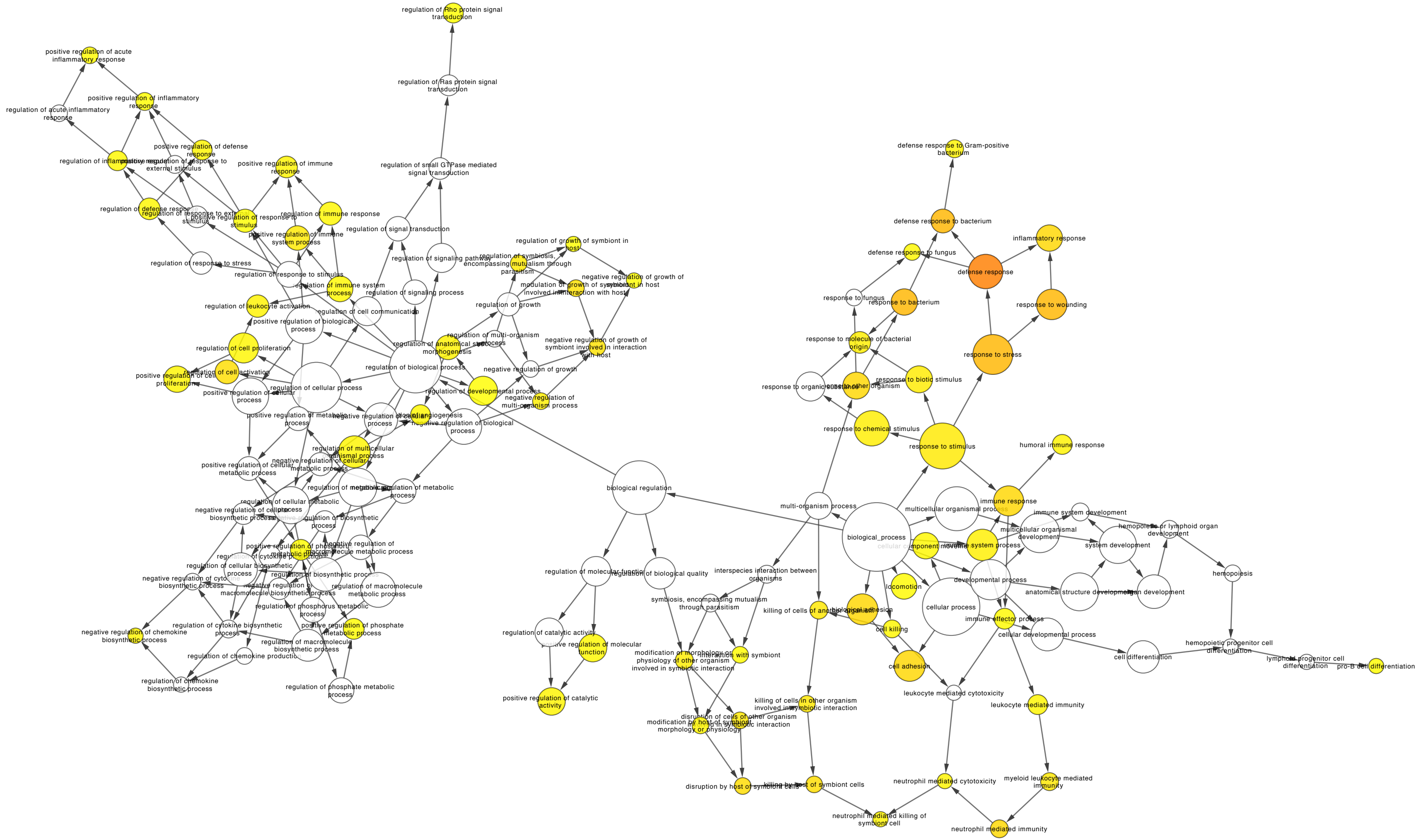
